# Genome scale CRISPR screens identify actin capping proteins as key modulators of therapeutic responses to radiation and immunotherapy

**DOI:** 10.1101/2024.01.14.575614

**Authors:** Shan Xin, Paul A. Renauer, Chuanpeng Dong, Yanzhi Feng, Ping Ren, Qianqian Lin, Feifei Zhang, Narges Tafreshi, Peter M. Glazer, Sidi Chen, Nipun Verma

## Abstract

Radiotherapy (RT), is a fundamental treatment for malignant tumors and is used in over half of cancer patients. As radiation can promote anti-tumor immune effects, a promising therapeutic strategy is to combine radiation with immune checkpoint inhibitors (ICIs). However, the genetic determinants that impact therapeutic response in the context of combination therapy with radiation and ICI have not been systematically investigated. To unbiasedly identify the tumor intrinsic genetic factors governing such responses, we performed a set of genome-scale CRISPR screens in melanoma cells for cancer survival in response to low-dose genotoxic radiation treatment, in the context of CD8 T cell co-culture and with anti-PD1 checkpoint blockade antibody. Two actin capping proteins, *Capza3* and *Capg*, emerged as top hits that upon inactivation promote the survival of melanoma cells in such settings. *Capza3* and *Capg* knockouts (KOs) in mouse and human cancer cells display persistent DNA damage due to impaired homology directed repair (HDR); along with increased radiation, chemotherapy, and DNA repair inhibitor sensitivity. However, when cancer cells with these genes inactivated were exposed to sublethal radiation, the persistence of DNA damage promoted activation of the STING pathway, induction of inhibitory *CEACAM1* ligand expression and resistance to CD8 T cell killing. Patient cancer genomics analysis reveals an increased mutational burden in patients with inactivating mutations in *CAPG* and/or *CAPZA3*, at levels comparable to other HDR associated genes. There is also a positive correlation between *CAPG* expression, CD8 T cell tumor infiltration and response to immunotherapy among patient datasets. Our results unveil the critical roles of actin binding proteins for efficient HDR within cancer cells and demonstrate a previously unrecognized regulatory mechanism of therapeutic response to radiation and immunotherapy.

## Introduction

Radiotherapy (RT), is a fundamental treatment for malignant tumors and is used for curative and palliative purposes in over half of cancer patients ^1,2^. The primary mechanism of action for radiation is the production of DNA double strand breaks (DSBs). As a response to DSBs, a robust DNA damage response is elicited that coordinates DNA damage detection and DNA repair, which is primarily through non-homologous end-joining (NHEJ) or homology-directed repair (HDR) ^3–5^. The dramatic increase in DSBs within cancer cells following genotoxic therapy like radiation can overwhelm this repair machinery and induce cell cycle arrest and cell death ^6^.

Over the past decade immune checkpoint inhibitors (ICI) have shown remarkable efficacy against a variety of cancers; however, many patients are unresponsive, fail to achieve a complete response or suffer frequent relapses ^7,8^. One promising strategy to increase the efficacy of ICI is to combine them with conventional therapies such as radiation. Prior studies have shown that radiation can trigger the immunogenic cell death of cancer cells and lead to the release of tumor-associated antigens ^9,10^, remodeling of the tumor microenvironment and activation of local immune cells ^11–14^, and the triggering of a systemic anti-tumor response ^15–18^. Although these studies suggest a possible synergism between radiation and ICI the global landscape of genetic determinants underlying the responses to the combination of radiation and immunotherapy has not been systematically explored before.

Here, we conducted a set of genome-scale CRISPR screens in the context of radiation plus immunotherapy. Our screen in B16F10 mouse melanoma cells identified actin capping proteins, *Capza3* and *Capg*, as important regulators that modulate cancer cell survival in the setting of low-dose genotoxic radiation, CD8 T cell co-culture and anti-PD1 antibody treatment. We also unexpectedly found that inactivation of CAPZA3 and CAPG proteins led to increased DNA damage after radiation treatment due to impaired HDR. Further investigation revealed the link between these actin capping proteins to the induction of the STING pathway and expression of T-cell inhibitory ligands and illustrated how HDR deficiency and chronic STING activation can promote resistance to CD8 T cell cytotoxicity.

## Results

### Genome-scale CRISPR screens identified genetic regulators of cancer cell survival in response to radiation and ICI

To systematically identify genes that regulate cancer cell response to radiation in combination with immunotherapy, we set out to perform a set of genome-scale CRISPR screens (**Figure 1a**). We used the classical B16F10 mouse melanoma cell line and transduced it with lentiviral vectors to constitutively express Cas9 and a model antigen ovalbumin (OVA), generating B16F10-OVA-Cas9 cells (hereafter referred to as B16F10 cells) (**Figure S1a**). We then created a pool of mutagenized B16F10 cells using a genome-scale guide RNA (gRNA) lentiviral library (Brie) ^19^, with a MOI of 0.3 to ensure that each cell harbored a single gRNA. We use CD8 T cells isolated from OT-I transgenic mice, which contain an engineered T-cell receptor (TCR) specific for the OVA antigen. These CD8 T were activated for 3 days with OVA peptide, resulting in upregulation of both activation and exhaustion markers (**Figure S1b**). Prior to performing the screen, we tested the editing efficiency and assessed the impact of specific gene knockout (KO) on B16F10 cell survival. Using a gRNA targeting the *Pdl1* locus (encoding the mouse PD-L1), we confirmed effective gene disruption (**Figure S1c**). Notably, *Pdl1* KO cells showed increased apoptosis following co-culture with OT-I CD8 T cells (**Figure S1d**).

**Figure 1.**
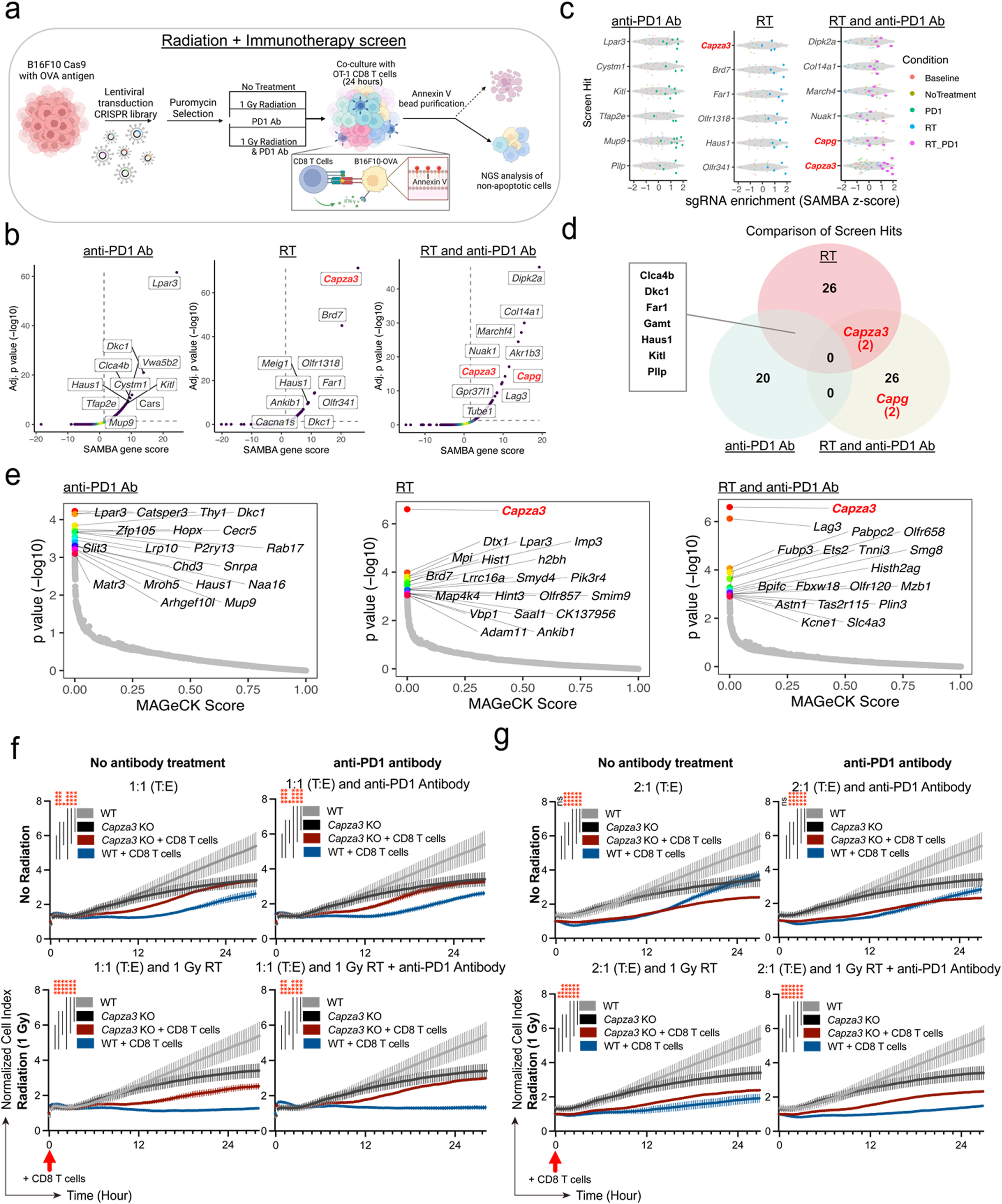
Genome-wide B16F10 CRISPR screens reveal that inactivation of actin capping proteins, *Capg* and *Capza3*, promotes B16F10 melanoma cell survival following radiation and co-culture with anti-PD1 antibody and CD8 cytotoxic T cells. **a,** Schematic of the B16F10 genome-scale CRISPR KO screens. **b,** Scatter plots for SAMBA screen analyses from surviving B16F10 following co-culture with CD8 cytotoxic T cells. Cells were co-cultured with anti-PD1 antibody (left), following treatment with 1 Gy low-dose radiation (middle) and following treatment with 1 Gy low-dose radiation and anti-PD1 antibody (right). Results shown are from 2 biologic repeats of the screen. **c,** Top 6 gRNAs enriched in surviving B16F10 cells following co-culture with CD8 cytotoxic T cells. Cells were co-cultured with anti-PD1 antibody (left), following treatment with 1 Gy low-dose radiation (middle) and following treatment with 1 Gy low-dose radiation and anti-PD1 antibody (right). Results shown are from 2 biologic repeats of the screen. **d,** Venn diagram of enriched targets (defined as q-value < 0.05) in different screen conditions. Number of gRNAs enriched for *Capza3* and *Capg* targets are shown in parentheses. **e,** Scatter plots for MAGeCK-RRA screen analyses from surviving B16F10 following co-culture with CD8 cytotoxic T cells. Cells were co-cultured with anti-PD1 antibody (left), following treatment with 1 Gy low-dose radiation (middle) and following treatment with 1 Gy low-dose radiation and anti-PD1 antibody (right). Results shown are from 2 biologic repeats of the screen. **f,** Cell attachment index of WT B16F10 and *Capza3* KO B16F10 co-cultured with OT-1 T cells. Cells were either just exposed to OT-1 CD8 T cells (1:1 T:E) or exposed to CD8 T cells with: anti-PD1 antibody (1:1 T:E and anti-PD1 Antibody), 1 Gy low-dose radiation (1:1 T:E and 1 Gy RT) or 1 Gy low-dose radiation and anti-PD1 antibody (1:1 T:E and 1 Gy RT + anti-PD1 Antibody). Experiments used a 1:1 T:E (tumor: effector) ratio. Time point of addition of OT-1 CD8 T cells is shown by an arrow. Data shown as means ± SEM. *P* value was determined by one-way ANOVA. Data are representative of three biological replicates. **g,** Cell attachment index of WT B16F10 and *Capza3* KO B16F10 co-cultured with OT-1 T cells. Cells were either just exposed to OT-1 CD8 T cells (2:1 T:E) or exposed to CD8 T cells with: anti-PD1 antibody (2:1 T:E and anti-PD1 Antibody), 1 Gy low-dose radiation (2:1 T:E and 1 Gy RT) or 1 Gy low-dose radiation and anti-PD1 antibody (2:1 T:E and 1 Gy RT + anti-PD1 Antibody). Experiments used a 2:1 T:E (tumor: effector) ratio. Time point of addition of OT-1 CD8 T cells is shown by an arrow. Data shown as means ± SEM. *P* value was determined by one-way ANOVA. Data are representative of three biological replicates.

To identify factors that influence CD8 T cell killing of B16F10 cells under different therapeutic conditions, we set up four treatment conditions in this set of screens: (1) no treatment (co-culture with OT-I CD8 T cells only), (2) low dose radiation (1 Gray (Gy)) prior to co-culture, (3) anti-PD1 antibody during co-culture, and (4) the combination of low dose radiation (1 Gy) prior to co-culture and anti-PD1 antibody during co-culture (**Figure 1a)**. We then collected surviving B16F10 mutant cell populations following 1 day of co-culture along with the baseline control of B16F10 cells transduced with the Brie CRISPR library but prior to any treatment (radiation, co-culture, or anti-PD1). We performed Annexin V-based column purification to enrich non-apoptotic cells for subsequent next-generation sequencing (NGS) of the gRNA cassette (**Figure S2a-b**).

Screen data processing and quality metrics showed robust screen performance (**Dataset S1**): (1) the screen replicates clustered with each other and away from the baseline controls (**Figure S2c**); (2) full coverage (near 100%) of the library was detected in all samples (**Figure S2d**); and (3) principal component analysis (PCA) showed the divergence of RT, PD-1, and combination treatment groups from the control groups in the PCA map (**Figure S2e**). We analyzed gRNA enrichment and depletion using a generalized linear model based statistical analysis (SAMBA), as well as a classical MAGECK analysis (**Methods**) ^20^. SAMBA analysis identified a set of enriched gRNAs and genes under each condition, including 35, 27, and 28 genes in RT, PD-1, and RT+PD1 treated cells, respectively (**Dataset S1**). The screen analysis revealed that *Capza3* was the topmost enriched gene in the RT screen group; followed by *Brd7,* a gene encoding a key component of the SWI-SNF complex and an epigenetic regulator ^21,22^. Interestingly, *Capza3* was also an enriched gene in the RT plus PD1 screen condition (**Figure 1b-d)**. Other notable genes enriched in RT+PD1 included *Dipk2a* (encoding Divergent Protein Kinase Domain 2A), *March4* (encoding a member of the MARCH family of membrane-bound E3 ubiquitin ligases), and *Lag3* (encoding a canonical immune checkpoint protein) (**Figure 1b-d)**. The PD1 screen condition’s top enriched gene was *Lpar3* (encoding Lysophosphatidic Acid Receptor 3, a G protein-coupled receptor family member) (**Figure 1b-d)**. Analysis using an independent MAGeCK analysis algorithm also confirmed that *Capza3* was the top enriched gene in both RT alone as well as RT plus PD-1 screen conditions (**Figure 1e; Dataset S1**).

The actin capping proteins encoding genes *Capza3* and *Capg* were significantly enriched among surviving B16F10 cells in the RT and RT+PD-1 condition (for *Capza3*) or in the RT+PD1 condition alone (for *Capg*), but neither was enriched in the PD1 alone condition (**Figure 1b-d**), hinting that these two targets may modulate the effect of radiation response in B16F10 cells (expressing OVA) in co-culture with CD8 T cells (OT-I), with or without PD1 antibody treatment. Given the prominence of *Capza3* in the screen, we prioritized it for further investigation through targeted gene studies. We used 2 independent *Capza3* gRNAs enriched in the screen to perform individual gene KO by lentiviral transduction into B16F10 cells and found that both result in high efficiency editing of the *Capza3* gene through T7E1 assay and next generation sequencing (**Figure S3a-b**), and cause reduction of *Capza3* at the mRNA (**Figure S3c**) and protein levels (**Figure S3d**). We confirmed that *Capza3* KO B16F10 cells transduced with either *Capza3* gRNA 2 or gRNA 3 had similar expression and presentation of the OVA antigen and the PD-L1 ligand as WT cells transduced with a non-targeting gRNA (**Figure S3e**).

To test the effect of *Capza3* KO on cell survival, we generated *Capza3* KO cells and used real-time cell analysis (RTCA) assay to confirm the enhanced survival of *Capza3* KO cells in co-culture with OT-I T cells under both RT and RT+PD-1 conditions. This survival benefit was observed at tumor-to-effector (T:E) ratios of 1:1 (**Figure 1f**) and 2:1 (**Figure 1g**). Intriguingly, despite their survival advantage under co-culture condition with CD8 T cells, *Capza3* KO cells cultured without CD8 T cells exhibited impaired proliferation *in-vitro* compared to WT controls (**Figure 1f-g**).

### B16F10 *Capza3* KO cells show increased DNA damage due to impaired homology-directed repair

Given enrichment of *Capza3* gRNA in radiation-containing screen conditions (RT or RT+PD-1) and the role of actin in DNA DSB repair ^23^, we hypothesized that *Capza3* may play a role in DNA damage response and maintaining genomic stability. To test this, we decided to investigate further whether inactivation of *Capza3* impacted DNA damage repair. Interestingly, although the *Capza3* gRNAs were enriched in the surviving tumor cells following co-culture, we found that *Capza3* KO B16F10 cells showed increased persistence of DNA damage markers (53BP1, γH2AX and pRPA) following radiation treatment compared to WT cells (**Figure 2a-d**) ^24–26^. The difference between WT and KO cells was more pronounced at low dose radiation (1 Gy) but was also significant with higher doses (5 Gy and 10 Gy) (**Figure 2d**). In addition, when normalized to the control no-treatment condition *Capza3* KO cells showed reduced colony forming ability than WT cells after exposure to radiation (**Figure 2e**).

**Figure 2.**
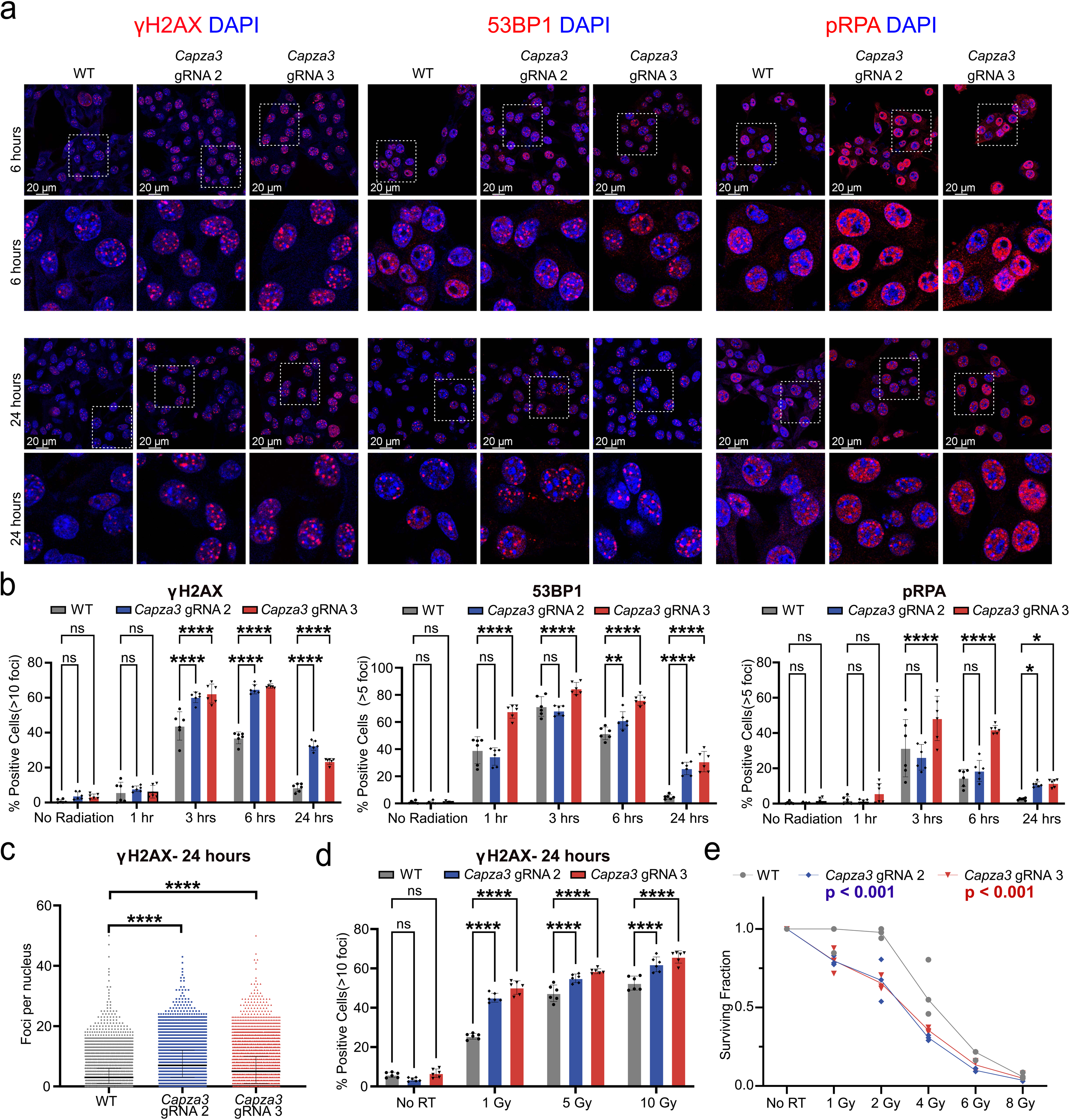
B16F10 *Capza3* KO cells show increased DNA damage after exposure to radiation. **a,** Immunofluorescence staining of γH2AX (left), 53BP1 (middle) and phospho-RPA2 (pRPA, right) foci in WT and *Capza3* KO cells at 6 hours and 24 hours after low-dose (1 Gy) radiation treatment. Representative nuclei from each image are enlarged. **b,** Quantification of γH2AX (left), 53BP1 (middle) and pRPA (right) foci in WT and *Capza3* KO cells at different timepoints following low-dose (1 Gy) radiation treatment. Significance testing was performed with two-way ANOVA, three biological replicates were done for each sample and treatment condition. **c,** Quantification of total γH2AX foci in individual WT and *Capza3* KO cells 24 hours after low-dose (1 Gy) radiation treatment. Significance testing was performed with one-way ANOVA, individual cell data from three biological replicates were pooled together for each sample. **d,** Quantification of γH2AX foci in WT and *Capza3* KO cells 24 hours after exposure to different doses of radiation treatment. Significance testing was performed with two-way ANOVA, three biological replicates were done for every sample and treatment condition. **e,** Colony forming ability (CFA) of WT and *Capza3* KO cells after exposure to different doses of radiation treatment. The survival fraction for each treatment was determined after normalization with the colony number seen in the no radiation treatment control for each cell line. Significance testing was performed with two-way ANOVA. Significance for both WT vs *Capza3* gRNA 2 and WT vs *Capza3* gRNA 3 is shown. Three biological replicates were done for each sample and treatment condition. For all experiments, WT cells were transduced with a lentiviral vector expressing a non-targeting control gRNA. * p < 0.05, ** p < 0.01, *** p < 0.001, **** p < 0.0001.

We found that *Capza3* KO cells had reduced RAD51 foci, a central homologous repair protein, at 6 hours after radiation treatment (**Figure 3a**). Furthermore, using HDR and NHEJ extrachromosomal luciferase reporters ^27^ (**Figure 3b**) we found that *Capza3* KO cells showed a marked reduction in HDR efficiency (**Figure 3c**). As an actin capping protein we predicted that loss of *Capza3* might impair DNA repair through disrupted actin remodeling. To test this, we inhibited the Arp2/3 complex, another actin regulator, using CK-666 (**Figure 3d-e**). WT and *Capza3* KO cells were cultured for 1 day at different concentrations of an Arp2/3 inhibitor CK-666. Cells were then exposed to radiation and γH2AX staining was done 24 hours after radiation treatment. Although Capza3 KO cells exhibited higher levels of DNA damage foci after radiation compared to WT cells prior to CK-666 treatment, WT cells cultured with increasing doses of CK-666 showed increasing DNA damage foci after radiation treatment, whereas *Capza3* KO cells displayed minimal changes (**Figure 3d**). Similarly, at baseline *Capza3* KO cells have reduced HDR efficiency compared to WT cells. With CK-666 treatment HDR efficiency in WT cells was significantly decreased whereas HDR efficiency in *Capza3* KO cells remained unaffected (**Figure 3e**). Next, we used laser micro-irradiation to observe the movement of DNA damage γH2AX foci following subnuclear-targeted radiation. Following laser micro-irradiation, *Capza3* KO cells showed increased concentration of γH2AX foci at the irradiated stripe and nuclear periphery, whereas WT cells showed γH2AX foci distributed throughout the nucleus (**Figure 3f**, **Figure S4a-d**). We also used laser-micro-irradiation to target regions of heterochromatin (identified as Hoechst bright DNA within the nucleus), euchromatin (identified as DNA within the nucleolus) and the nuclear membrane. Notably *Capza3* KO showed a different pattern in the movement of damaged heterochromatin DNA compared to WT cells, with increased concentration of DNA damage at the nuclear periphery in *Capza3* KO cells at 2 hours following laser micro-irradiation (**Figure 3g**). Finally, to test whether this persistent DNA damage could promote micronuclei formation, we used both Hoechst staining as well as an H2B fluorescent reporter to monitor micronuclei formation following radiation treatment. 24-hour live cell imaging of WT and *Capza*3 KO cells showed increased micronuclei formation in the KO cells following radiation treatment (**Figure 3h**).

**Figure 3.**
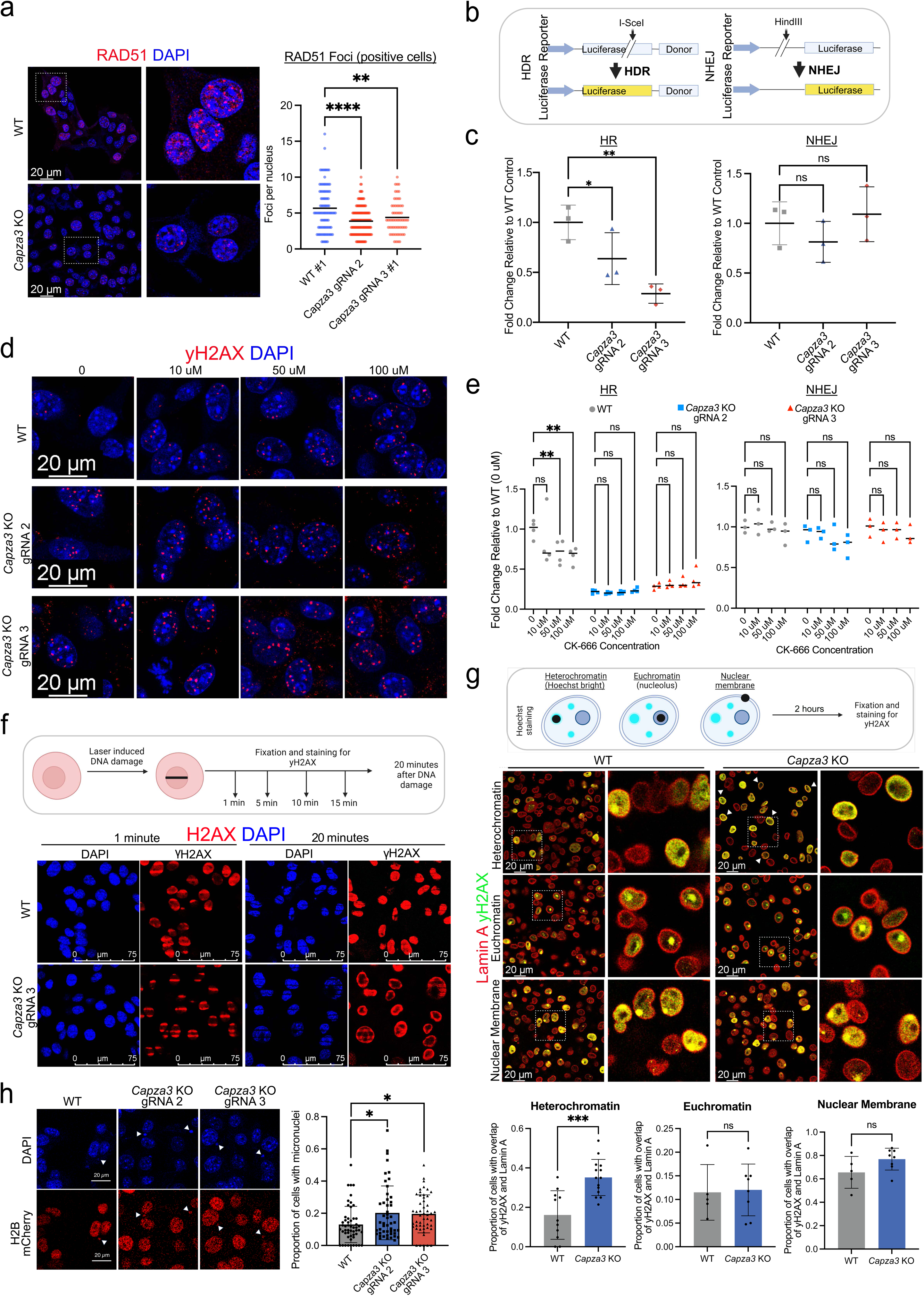
B16F10 *Capza3* KO cells have impaired HDR compared to WT cells. **a,** Immunofluorescence staining of RAD51 foci in WT and *Capza3* KO cells at 6 hours after low-dose (1 Gy) radiation treatment. Representative nuclei are enlarged and presented to the right. Quantification of the RAD51 foci per nucleus in RAD51 positive cells is shown to the right. Significance testing was performed with one-way ANOVA, three biological replicates were done for every sample and treatment condition. **b,** Schematic of extra-chromosomal reporters used to analyze efficiency of HDR and NHEJ in WT and *Capza3* KO B16F10 cells. **c,** Quantification of luciferase activity, compared to WT cells, 48 hours after transfection with linearized HDR (left) and NHEJ (right) extrachromosomal reporters. Significance testing was performed with two-way ANOVA. Three biological replicates were done for each sample. **d,** Representative immunofluorescence staining of γH2AX foci in WT and *Capza3* KO cells at 24 hours after 1 Gy radiation treatment following incubation with varying concentrations of CK-666. **e,** Quantification of luciferase activity, compared to untreated WT cells, 48 hours after transfection with linearized HDR (left) and NHEJ (right) extrachromosomal reporters. Significance testing was performed with two-way ANOVA, 3-4 biological replicates were done for each sample. **f,** Schematic of laser micro-irradiation experiment to induce DNA damage in an irradiated stripe followed by fixation and staining with γH2AX at different time points. Representative immunofluorescence images of γH2AX DNA damage foci 1 minute and 20 minutes following laser micro-irradiation in WT and *Capza3* KO cells are shown. **g,** Schematic of laser micro-irradiation to induce DNA damage at heterochromatin (Hoechst bright), euchromatin (nucleolus) and nuclear membrane regions followed by fixation and staining with γH2AX. Representative immunofluorescence images (top) and quantification (bottom) of γH2AX DNA damage foci 2 hours following laser micro-irradiation in WT and *Capza3* KO cells are shown. **h,** 24 hours live cell imaging of WT and *Capza3* KO cells with Hoechst staining and a H2B fluorescent reporter to monitor proportion of cells that show micronuclei formation following 1 Gy radiation treatment. Micronuclei formation was quantified in approximately 50 representative fields of view. Cells with at least one micronuclei were considered positive. Significance testing was performed with one-way ANOVA, two biological replicates were done, with 20-30 fields quantified for each replicate. For all experiments, WT cells were transduced with a lentiviral vector expressing a NTC gRNA. * p < 0.05, ** p < 0.01, *** p < 0.001, **** p < 0.0001.

### *Capza3* KO cells show enhanced *STING* pathway activation and increased expression of the immunosuppressive ligand *Ceacam1*

Our data above showed that *Capza3* inactivation leads to reduced HDR efficiency and persistent DNA damage following radiation treatment. However, the mechanism underlying the increased resistance of *Capza3* KO cells to CD8 T cell killing remained unclear. To further investigate this, we performed transcriptome profiling using RNA-seq in WT and *Capza3* KO B16F10 cells. Differential expression (DE) analysis of RNA-seq data (**Methods**) revealed transcriptomic differences between WT and *Capza3* KO cells, and between RT and untreated samples (**Figure S5a-b**). We verified that replicates of the same genotype showed higher correlation as compared to between genotypes/groups, and clustered together on PCA (**Figure S5c-d**). Genes encoding transcription factors associated with the STING pathway (*Irf1, Irf9,* and *Stat2*) showed increased activity in *Capza3* KOs compared to WT cells both before and after radiation treatment (**Figure 4a**, **Figure S5e**). Activation of the STING pathway leads to increased expression of type I interferons, signaling via IRF9 and STAT2, expression of IRF1 and ultimately increased expression of IRF1 target genes (**Figure 4b**). Further analysis of the IRF1 targets showing greatest change in expression between WT and *Capza3* KOs revealed increased expression of *Ceacam1*, a known target of IRF1 ^28–30^, in *Capza3* KO cells (**Figure 4c**).

**Figure 4.**
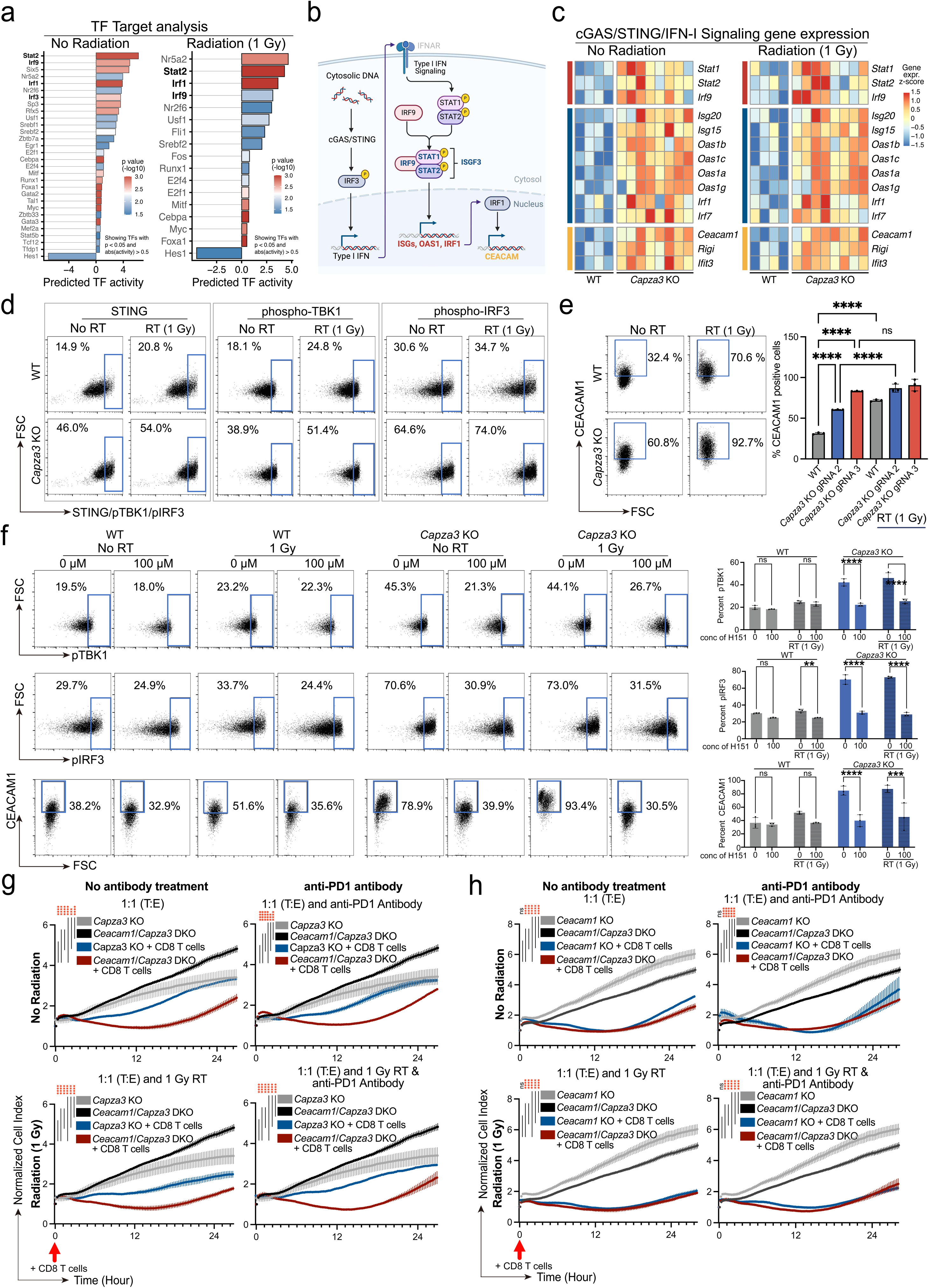
B16F10 *Capza3* KO cells show increased activation of the STING pathway and increased expression of *Ceacam1* compared to WT cells and inactivation of *Ceacam1* in *Capza3* KO cells removes their survival advantage following radiation treatment and co-culture with CD8 T cells. **a,** Analysis of transcription factor pathways that are significantly upregulated in *Capza3* KO compared to WT B16F10 cells before (left) and after (right) low-dose (1 Gy) radiation treatment. Transcription factors with p-value <0.05 and absolute activity.> 0.5 are shown. Analysis was done on bulk RNA-Seq data of WT and *Capza3* KO B16F10 cells. **b,** Schematic of cGAS/STING activation pathway with downstream expression of IRF1 and IRF1 target genes. **c,** Targets of the IRF1 transcription factor that showed the greatest upregulation in *Capza3* KO compared to WT B16F10 cells before (left) and after (right) low-dose (1 Gy) radiation treatment. **d,** Representative flow plots of STING pathway staining (STING, phospho-TBK1, phospho-IRF3) in WT and *Capza3* KO B16F10 cell lines before and after low-dose (1 Gy) radiation treatment. **e,** Representative flow plots of CEACAM1 staining in WT and *Capza3* KO B16F10 cell lines before and after low-dose (1 Gy) radiation treatment. Quantification of CEACAM1 positive cells is shown on the right. Significance testing was performed with two-way ANOVA, three biological replicates were done for each sample and treatment condition. **f,** Representative flow plots of STING pathway staining (phosphor-TBK1, phosphor-IRF3) in WT and *Capza3* KO B16F10 cell lines with and without pre-treatment with STING inhibitor H151 and before and after low-dose (1 Gy) radiation treatment. Representative plots are showed on the left and quantification of positive cells is shown on the right for phospho-TBK1 (top), phospho-IRF3 (middle) and CEACAM1 (bottom). Significance testing was performed with two-way ANOVA, three biological replicates were done for each sample and treatment condition. **g,** Cell attachment index of *Capza3* KO B16F10 and *Capza3*/*Ceacam1* DKO B16F10 cells co-cultured with OT-1 T cells. Cells were either just exposed to OT-1 CD8 T cells (1:1 T:E) or exposed to CD8 T cells with: 1 Gy low-dose radiation (1:1 T:E and 1 Gy RT), anti-PD1 antibody (1:1 T:E and anti-PD1 Antibody) or 1 Gy low-dose radiation and anti-PD1 antibody (1:1 T:E and 1 Gy RT + anti-PD1 Antibody). Experiments used a 1:1 T:E (tumor: effector) ratio. Time point of addition of OT-1 CD8 T cells is shown by an arrow. Data shown as means ± SEM. *P* value was determined by two-way ANOVA. Data are representative of three biological replicates. **h,** Cell attachment index of *Ceacam1* KO B16F10 and *Capza3*/*Ceacam1* DKO B16F10 cells co-cultured with OT-1 T cells. Cells were either just exposed to OT-1 CD8 T cells (1:1 T:E) or exposed to CD8 T cells with: 1 Gy low-dose radiation (1:1 T:E and 1 Gy RT), anti-PD1 antibody (1:1 T:E and anti-PD1 Antibody) or 1 Gy low-dose radiation and anti-PD1 antibody (1:1 T:E and 1 Gy RT + anti-PD1 Antibody). Experiments used a 1:1 T:E (tumor: effector) ratio. Time point of addition of OT-1 CD8 T cells is shown by an arrow. Data shown as means ± SEM. *P* value was determined by two-way ANOVA. Data are representative of three biological replicates. For RNA-Seq analysis, n = 2 biologic replicates for each cell line (WT #1, WT #2, *Capza3* gRNA 2 KO #1, *Capza3* gRNA 2 KO #2, *Capza3* gRNA 3 KO #1, *Capza3* gRNA 3 KO #2) and each condition (no radiation or after 1 Gy radiation) for a total of 24 samples were submitted for sequencing. * p < 0.05, ** p < 0.01, *** p < 0.001, **** p < 0.0001.

As *Ceacam1* is a known CD8 T cell inhibitor ligand, we hypothesized that enhanced activation of the STING pathway and increased downstream expression of CEACAM1 in *Capza3* KO cells may explain their survival advantage following co-culture with CD8 T cells and decided to explore this target further. We confirmed that compared to WT cells, *Capza3* KO cells show greater activation of the STING pathway and increased expression of CEACAM1 by flow cytometry of phospho-TBK1, phospho-IRF3 and CEACAM1. Moreover, CEACAM1 expression is further increased in both WT and *Capza3* KO cells following radiation treatment (**Figure 4d-e**). Importantly, treatment of *Capza3* KO cells with H151, a STING inhibitor, reduced activation of the STING pathway and reduced induction of CEACAM1 expression following radiation treatment (**Figure 4f**). Interestingly, impairing HDR in WT cells using the Arp2/3 inhibitor CK-666 also increased CEACAM1 expression in WT cells with and without radiation treatment (**Figure S6a**). Together, these findings suggest that CEACAM1 upregulation is a downstream response to STING pathway activation, triggered by persistent DNA damage and impaired HDR due to the loss of *Capza3*.

To determine whether increased CEACAM1 was responsible for the survival benefit of *Capza3* KO cells following radiation treatment and co-culture with CD8 T cells (**Figure 1f-g**), we inactivated *Ceacam1* in WT and *Capza3* KO B16F10 cells (**Figure S6b**). Using the RTCA assay, we found that inactivating *Ceacam1* in *Capza3* KO cells (double knockout or DKO) abrogated their survival advantage following co-culture with CD8 T cells (**Figure 4g-h, S6c-d**).

### *CAPG*/*CAPZA3* KO human cancer cells show impaired HDR, and increased CEACAM1 leads to a survival advantage when treated with radiation and anti-EGFR CAR-T

In *Capza3* KO B16F10 cells, persistent DNA damage and impaired HDR leads to activation of the STING pathway and upregulation of CEACAM1, promoting immune evasion. To determine if these findings are reproduced in human cells, we utilized human breast (MDA-MB-231) and pancreatic (PANC1) cancer cell lines that express the EGFR antigen. We generated *CAPG* (another actin capping protein enriched in the radiation and PD-1 treatment screen condition) and *CAPZA3* KO cells and confirmed inactivating mutations through next generation sequencing (**Figure S7a**). Multiple clonal *CAPG* and *CAPZA3* KO mutants in the MDA-MB-231 and PANC1 cell lines showed increased DNA damage following radiation treatment (**Figure 5a-c, S7b-c**) and reduced HDR repair rates, but no difference in NHEJ repair, using the extrachromosomal HDR and NHEJ luciferase reporters (**Figure 5d, S7d-e**).

**Figure 5.**
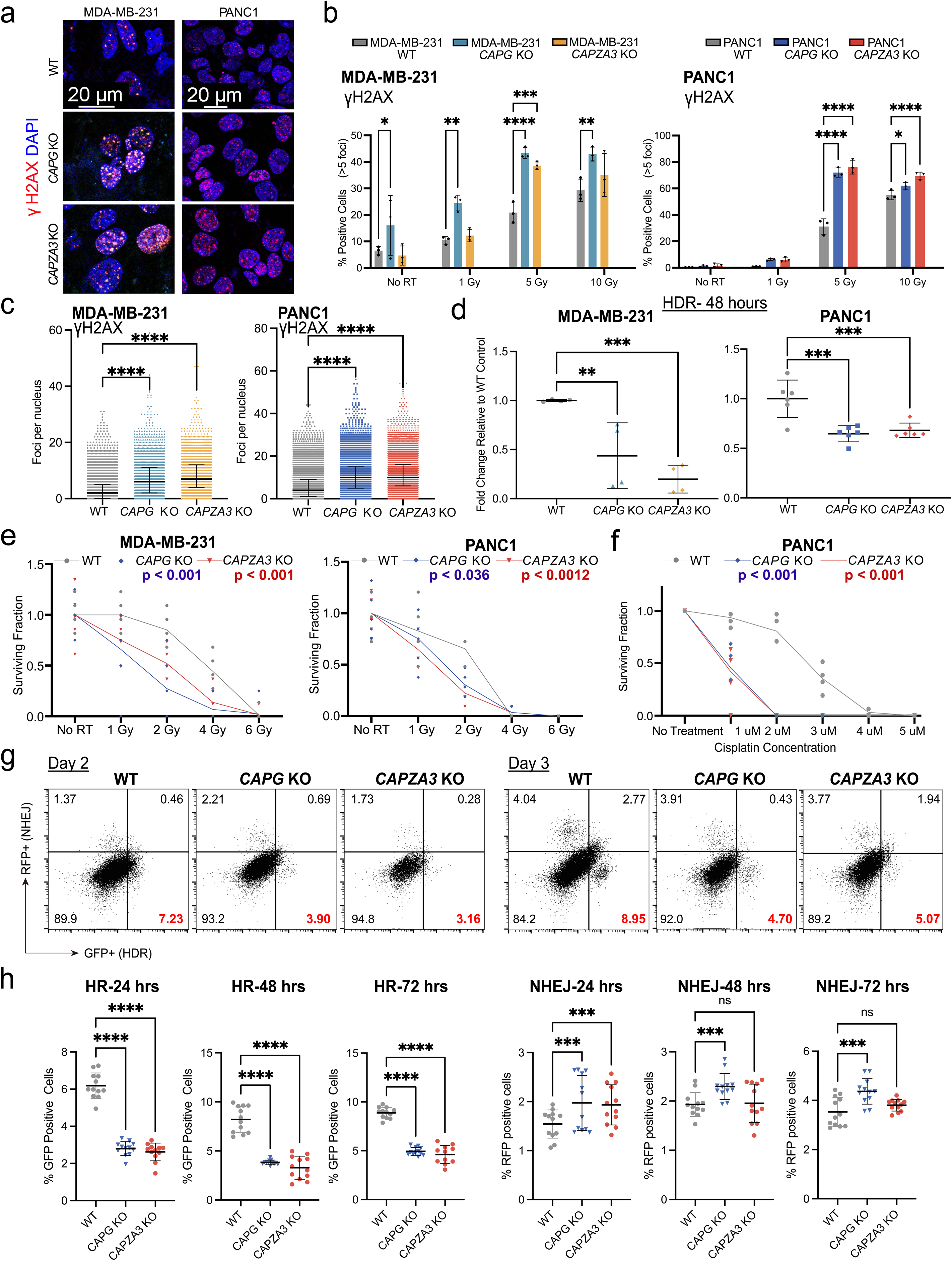
PANC1 and MDA-MB-231 *CAPG* and *CAPZA3* KO cell lines show increased DNA damage following radiation treatment and impaired HDR. **a,** Representative γH2AX immunofluorescence staining 24 hours after radiation treatment (5 Gy) in WT, *CAPG* and *CAPZA3* KO MDA-MB-231 and PANC1 cell lines. **b,** Quantification of γH2AX foci in WT, *CAPG* and *CAPZA3* KO MDA-MB-231 (left) and PANC1 (right) clonal cell lines 24 hours after different doses of radiation treatment. Significance testing was performed with two-way ANOVA, three biological replicates were done for each sample and treatment condition. **c,** Quantification of total γH2AX foci in individual cells in WT, *CAPG* and *CAPZA3* KO MDA-MB-231 (left) and PANC1 (right) clonal cell lines 24 hours after exposure to 5 Gy radiation. Significance testing was performed with one-way ANOVA, individual cell data from three biological replicates were pooled together for each sample. P-values are shown for comparisons of the KO line (*CAPG* KO or *CAPZA3* KO) to WT. **d,** Quantification of luciferase activity, compared to WT cells, 48 hours after transfection with a linearized HDR extrachromosomal reporter. Significance testing was performed with one-way ANOVA. 4 biological replicates were done for the MDA-MB-231 and 6 biological replicates were done for the PANC1. **e,** CFA of WT, *CAPG* and *CAPZA3* KO MDA-MB-231 (left) and PANC1 (right) cell lines after different doses of radiation treatment. Surviving fraction for each treatment was determined after normalization with the colony number seen in control condition without radiation treatment for each cell line. Significance testing was performed with two-way ANOVA, the p value for WT vs *CAPG* KO is shown in blue and the p value for WT vs *CAPZA3* KO is shown in red. Three biological replicates were done for each sample and treatment condition. **f,** CFA of WT, *CAPG* and *CAPZA3* KO PANC1 cell lines after treatment with different doses of cisplatin. Surviving fraction for each treatment was determined after normalization with the colony number seen in control condition without cisplatin treatment for each cell line. Significance testing was performed with two-way ANOVA, the p value for WT vs *CAPG* KO is shown in blue and the p value for WT vs *CAPZA3* KO is shown in red. Three biological replicates were done for each sample and treatment condition. **g,** Representative flow plots of RFP positive and GFP positive cells 48 hours and 72 hours after transfection of *Rosa26* targeting gRNA into PANC1 AAVS1 traffic light reporter (TLR) cells. **h,** Quantification of GFP positive cells (left, % cells with successful HDR) and RFP positive cells (right, % cells with successful NHEJ) 24, 48 and 72 hours after transfection of *Rosa26* gRNA in PANC1 *AAVS1* TLR WT, *CAPG* and *CAPZA3* KO clonal cell lines. Significance testing was performed with two-way ANOVA, four biological replicates were done for each sample. For all experiments, WT cells were passage-matched controls that underwent the procedure to generate KO cell lines (transfected with plasmid containing *CAPG* or *CAPZA3* targeting gRNA, sorted as single cells into a 96 well plate and expanded) but did not have a mutation at either the *CAPG* or *CAPZA3* locus. * p < 0.05, ** p< 0.01, *** p < 0.001, **** p < 0.0001.

Moreover, both MDA-MB-231 and PANC1 *CAPG* and *CAPZA3* KO cells showed reduced CFA after exposure to genotoxic agents such as radiation (**Figure 5e**) and cisplatin treatment (**Figure 5f**), suggesting sensitivity to DNA damage. To further verify the HDR defect seen in these KO cells, we generated a PANC1 intra-chromosomal HDR and NHEJ reporter line by integrating the DNA repair traffic light reporter (TLR) construct ^31^ into the *AAVS1* locus (PANC1-TLR). Using this line, we once again generated multiple *CAPG* and *CAPZA3* KO clonal lines. Following transfection with the *Rosa26* targeting gRNA we found that *CAPG* and *CAPZA3* KO lines showed lower HDR rates compared to WT cells (**Figure 5g-h, Figure S7f**).

We previously identified enhanced activation of the STING pathway in B16F10 cells through RNA-Seq analysis. Consistent with these findings, we confirmed increased STING and phospho-TBK1 protein levels in MDA-MB-231- and PANC1-*CAPG* and *CAPZA3* KO cells using flow cytometry (**Figure 6a-b, Figure S8a-b**). We also found increased *CEACAM1* gene expression at the mRNA level using qPCR (in MDA-MB-231 and PANC1 lines) (**Figure S8c)** and at the protein level in PANC1 cells using flow cytometry **(Figure 6c**). We then performed co-culture experiments using the EGFR expressing MDA-MB-231 and PANC1 lines and EGFR targeting CAR-T (**Figure 6d**, **S9a-c**). The EGFR CAR-T expressed both CEACAM1 and TIM3, which are known to have homophilic and heterophilic interactions with CEACAM1 on tumor cells, during maintenance culture (**Figure 6e**). We observed a survival advantage for MDA-MB-231 and PANC1 *CAPG* and *CAPZA3* KO lines compared to WT cells at low dose radiation (1 Gy) treatment prior to co-culture with EGFR CAR-T (**Figure 6f**).

**Figure 6.**
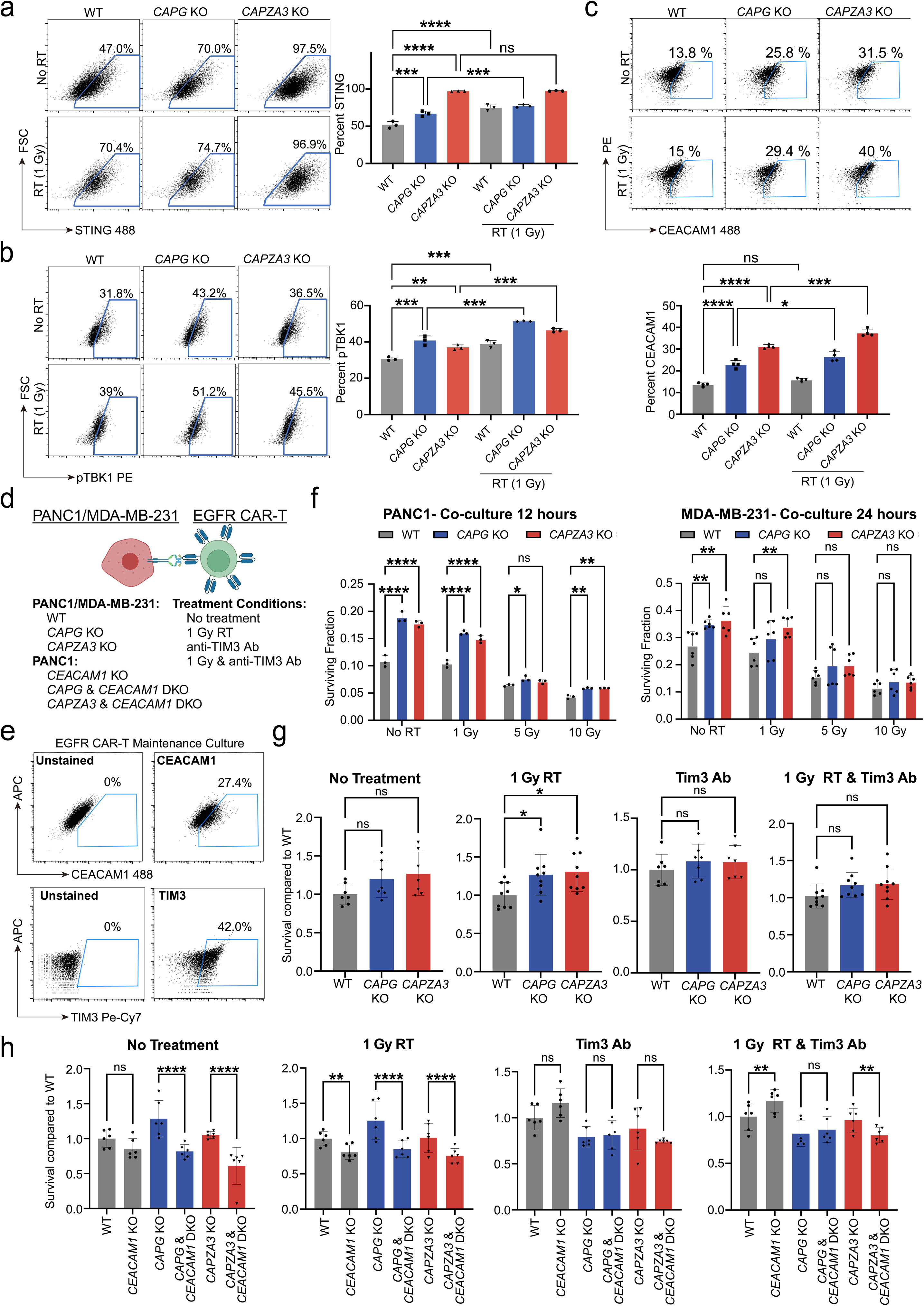
PANC1 *CAPG*/*CAPZA3* KO cells lose their survival advantage following co-culture with EGFR CAR-T after inactivation of *CEACAM1* or with co-culture with anti-TIM3 antibody. **a,** Representative flow plots of STING staining in WT, *CAPG* KO and *CAPZA3* KO PANC1 cell lines before and after low-dose radiation treatment (1 Gy). Representative plots are showed on the left and quantification of positive cells is shown on the right. Significance testing was performed with two-way ANOVA, three biological replicates were done for each sample and treatment condition. **b,** Representative flow plots of STING staining in WT, *CAPG* and *CAPZA3* KO PANC1 cell lines before and after low-dose (1 Gy) radiation treatment. Representative plots are showed on the left and quantification of positive cells is shown on the right. Significance testing was performed with two-way ANOVA, three biological replicates were done for each sample and treatment condition. **c,** Representative flow plots of CEACAM1-488 staining in WT, *CAPG* KO and *CAPZA3* KO PANC1 cell lines before and after low-dose (1 Gy) radiation treatment. Quantification of CEACAM1 positive cells is shown on the bottom panel. Significance testing was performed with two-way ANOVA, four biological replicates were done for each sample and treatment condition. **d,** Schematic of cell lines and treatment conditions used for co-culture experiments with PANC1 and MDA-MB-231 tumor cells and EGFR CAR-T. **e,** Representative flow plot of CEACAM1 and TIM3 flow analysis in EGFR CAR-T cells during regular maintenance culture. **f,** Analysis of cell survival after exposure to radiation and co-culture with EGFR CAR-T for WT, *CAPG* KO and *CAPZA3* KO PANC1 (left) and MDA-MB-231 (right) cell lines. Significance testing was performed with two-way ANOVA, three biological replicates were done for each sample and treatment condition. **g,** Cell survival of PANC1 *CAPG* and *CAPZA3* KO cell lines compared to WT cells following treatment with radiation and/or anti-TIM3 antibody and co-culture with EGFR CAR-T. Significance testing was performed with one-way ANOVA, three biological replicates were done for each sample. **h,** Cell survival of PANC1 KO cell lines (*CEACAM1*, *CAPG* and *CAPZA3* single and double KO) compared to WT cells following treatment with radiation and/or anti-TIM3 antibody and co-culture with EGFR CAR-T. Significance testing was performed with two-way ANOVA, three biological replicates were done for each sample and treatment condition. For all experiments, WT cells were passage-matched controls that underwent the procedure to generate KO cell lines (transfected with plasmid containing *CAPG* or *CAPZA3* targeting gRNA, sorted as single cells into a 96 well plate and expanded) but did not have a mutation at either the *CAPG* or *CAPZA3* locus. * p < 0.05, ** p< 0.01, *** p < 0.001, **** p < 0.0001.

To investigate whether increased expression of *CEACAM1* directly contributes to the survival advantage of *CAPZA3* KO human cancer cells, we performed coculture in the presence or absence of anti-TIM3 antibody. We found that the survival advantage of *CAPG*/*CAPZA3* KO PANC1 lines was lost if anti-TIM3 antibody was added during co-culture (**Figure 6g**). Furthermore, knocking out *CEACAM1* in *CAPG*/*CAPZA3* KO PANC1 lines also negated their survival advantage following co-culture with EGFR CAR-T (**Figure 6h**), suggesting that CEACAM1 is a direct mediator of resistance to CD8 T cell killing.

### *Capza3* KO B16F10 cells show increased activation of STING and expression of CEACAM1 in *in-vivo* tumor models

To validate our *in-vitro* findings we used *in-vivo* B16F10 mouse tumor models. First, we assessed tumor growth in immunocompromised nCR mice and immune competent C57BL/6 mice by subcutaneously injecting WT and *Capza3* KO cells and monitoring tumor growth over time (**Figure 7a**). We observed that *Capza3* KO tumors exhibited slower growth than WT tumors (**Figure 7b**), consistent with the reduced colony-formation (**Figure S10a**) and reduced proliferation (**Figure 1g-h**) we observed *in-vitro*. Next, we investigated *in-vivo* STING pathway activation and CEACAM1 expression by injecting WT and *Capza3* KO cells into nCR mice and exposing tumors to radiation (**Figure 7c**). Like our *in-vitro* studies, we found that *Capza3* KO tumors displayed increased phosphorylation of TBK1 and IRF3, along with elevated CEACAM1 expression, compared to WT tumors (**Figure 7d**). Radiation treatment further enhanced all three markers in both WT and KO tumors (**Figure 7d**), highlighting the increased activation of STING pathway and downstream CEACAM1 expression in *Capza3* KO tumors compared to WT tumors following radiation treatment.

**Figure 7.**
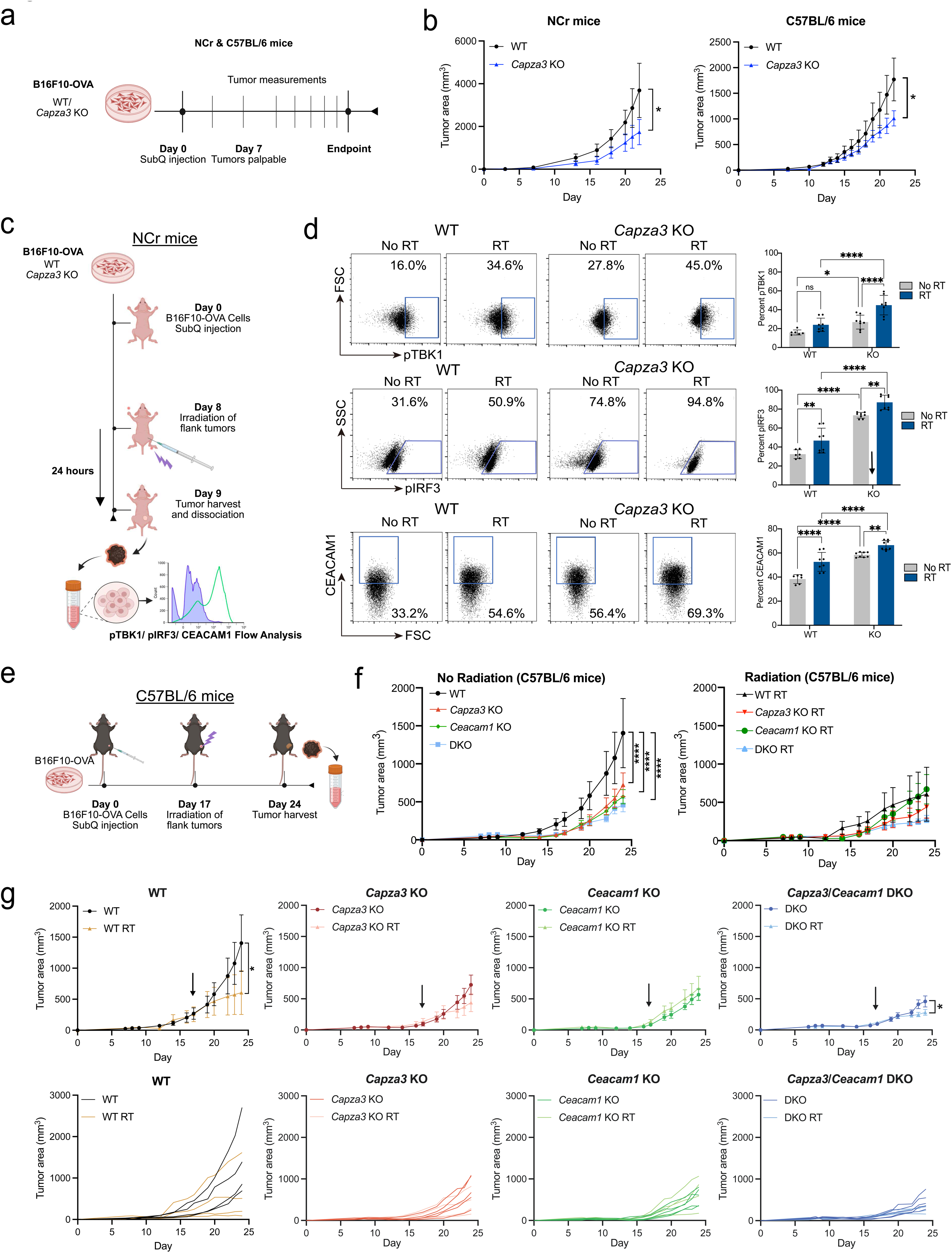
*Capza3* KO B16F10 cells show increased activation of STING and expression of CEACAM1 in *in-vivo* tumor models. **a,** Schematic of *in-vivo* tumor growth measurements in NCr and C57BL/6 mice. **b,** Tumor growth curves for WT and *Capza3* KO cells in NCr (left) and C57BL/6 mice. **c,** Schematic of *in-vivo* STING pathway and CEACAM1 analysis experiments using B16F10 cells in NCr mice. **d,** Representative flow plots of cGAS/STING pathway and CEACAM1 staining in WT and *Capza3* KO B16F10 tumors before and after 5 Gy radiation treatment in NCr mice: phospho-TBK1 (top), phospho-IRF3 (middle) and CEACAM1 (bottom). Quantification is shown on the right. Significance testing was performed with two-way ANOVA. A minimum of 3 biologic replicates were analyzed. **e,** Schematic of *in-vivo* tumor growth analysis of WT, *Capza3* KO, *Ceacam1* KO and *Capza3*/*Ceacam1* DKO B16F10 tumors in C57BL/6 mice. **f,** Tumor growth curves WT, *Capza3* KO, *Ceacam1* KO and *Capza3*/*Ceacam1* DKO B16F10 tumors with (right) and without (left) 5 Gy radiation treatment in C57BL/6 mice. Significance testing was performed with two-way ANOVA. A minimum of 3 biologic replicates were analyzed. **g,** Tumor growth curves of OVA expressing WT, *Capza3* KO, *Ceacam1* KO and *Capza3*/*Ceacam1* DKO B16F10 tumors with and without 5 Gy radiation treatment in C57BL/6 mice. Individual tumor growth curves for each biological replicate are shown below. Significance testing was performed with two-way ANOVA. A minimum of 3 biologic replicates were analyzed.

Next, we sought to examine tumor growth of WT, *Capza3* KO, *Ceacam1* KO and *Capza3*/*Ceacam1* DKO tumors in response to RT by injecting the cells into immune-competent C57BL/6 mice subcutaneously (**Figure 7e**). *Capza3* KO, *Ceacam1* KO, and *Capza3*/*Ceacam1* DKO tumors displayed significantly slower growth than WT tumors in the absence of radiation (**Figure 7f**, **Figure S10a**). We then assessed the differential response of each tumor genotype to radiation. While WT tumors exhibited a significant response to RT, *Capza3* KO tumors did not, suggesting a survival advantage for *Capza3* KO tumors under RT in the immunocompetent context, which aligns with our *in-vitro* findings (**Figure 7g**). Strikingly, *Capza3*/*Ceacam1* DKO tumors exhibited a robust response to RT, emphasizing the critical role of CEACAM1 downstream of *Capza3* KO in conferring a survival advantage against RT response within the C57BL/6 tumor model.

### *CAPZA3* and *CAPG* expression impacts tumor mutational burden, CD8 tumor cell infiltration, immunotherapy response and overall survival in cancer patient data sets

To evaluate the clinical relevance of our findings, we analyzed the TCGA dataset for *CAPZA3* and *CAPG* associated prognostic signatures of clinical outcome. Interestingly, we found that in-activating mutations for *CAPG* and *CAPZA3* frequently co-occur with mutations for other HDR associated genes (**Figure S11a**). Although we may expect that mutations of genes that function in the same pathway would be mutually exclusive, prior studies have shown that mutations in DNA repair genes frequently co-occur in numerous cancer types ^33^. Since there was co-occurrence of mutations in *CAPG/CAPZA3* and genes known to be associated with HDR, for the rest of our analyses we excluded any TCGA data samples that had a mutation in more than one HDR associated gene. Inactivating mutations in *CAPG/CAPZA3* led to increased tumor mutational burden and mutation counts, which was comparable to other HDR associated genes, and significantly increased compared to the unaltered control group of patients in a pan-cancer analysis (**Figure 8a, Figure S11b**). *CAPG* expression also showed a significant correlation with CD8 T cell infiltration (**Figure 8b**). Focusing on datasets in which patients with melanoma were treated with immune checkpoint inhibitors (ICI) we found that increased *CAPG* expression was associated with better response to ICI and improved survival (**Figure 8c**).

**Figure 8.**
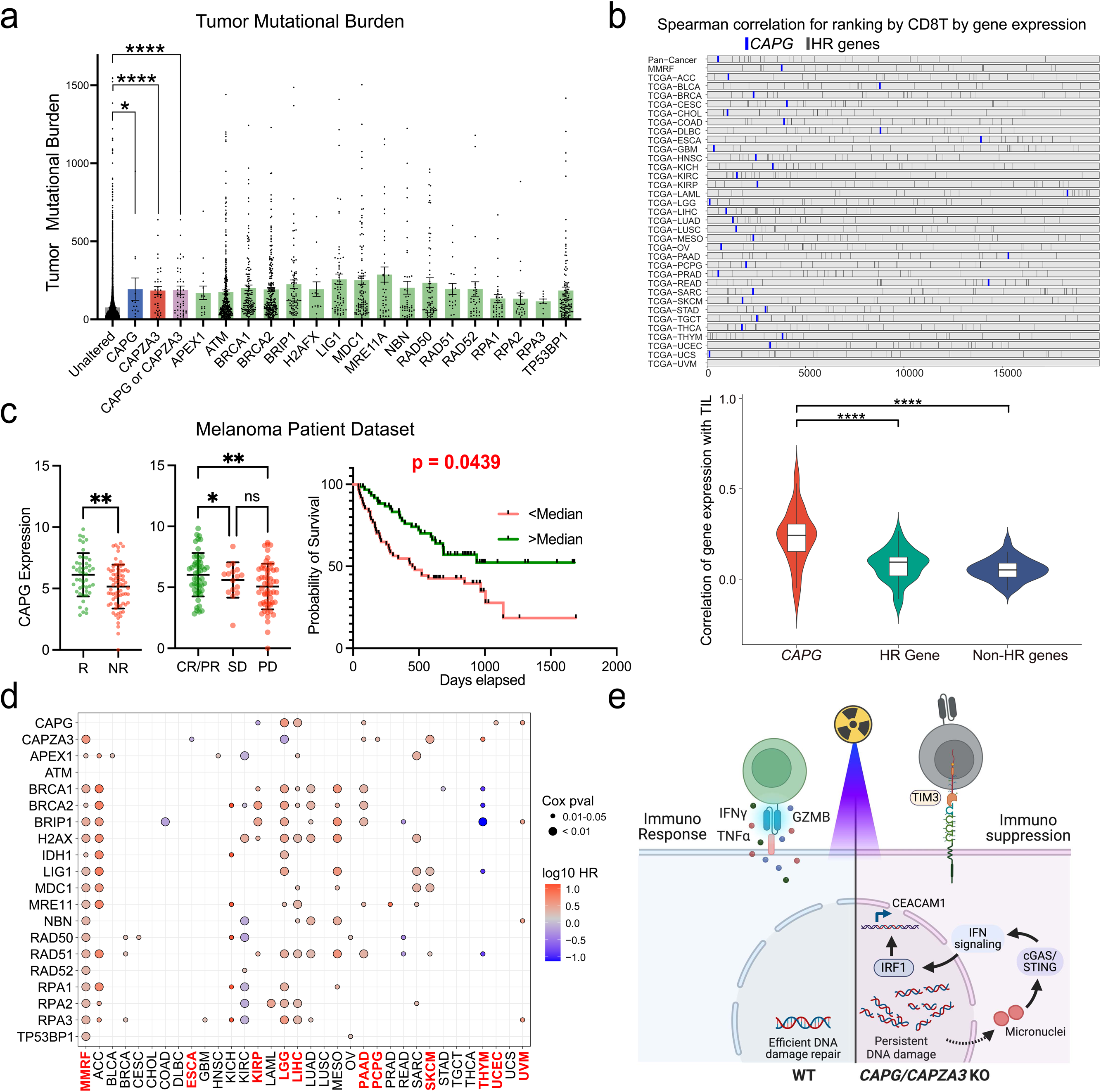
Patient dataset analysis of tumor mutational burden, CD8 T cell infiltration and overall survival based on expression of *CAPG*, *CAPZA3* and known HDR associated genes. **a,** Analysis of tumor mutational burden in patients with an isolated inactivating mutation in *CAPG* or *CAPZA3* or a mutation in a single HDR associated gene. P-values are shown for selected comparisons. Significance testing was performed with one-way ANOVA. **b,** Brick plot showing the correlation coefficient of CD8 T cell infiltration and HR gene expression compared to other genes in 33 TCGA cancer types. Each horizontal lane represents a TCGA cancer type, and the vertical lines indicate the ranking of HR genes. The correlation between CD8 T cell infiltration and the expression of each HR gene was measured using Spearman correlation. Correlation between gene expression and CD8 T cell infiltration for *CAPG*, all other HR genes and all non-HR genes is shown in the boxplot below. **c,** *CAPG* expression of melanoma cancer patients among responders (R) and non-responders (NR) to ICI therapy (left). *CAPG* expression of melanoma cancer patients among patients who showed complete or partial response (CR/PR), stable disease (SD) or progressive disease (PD) following ICI therapy based on RECIST criteria (middle). Significance testing was performed with one-way ANOVA. Survival curves for melanoma patients with *CAPG* expression above the median or below the median level, significance testing was performed with Log-rank (Mantel-Cox) test. **d,** Bubble plot illustrating the overall survival hazard ratios of *CAPG*, *CAPZA3* and other HR genes across 34 cancer types. Hazard ratios greater than 1, indicating unfavorable survival with higher expression, are colored red, while hazard ratios less than 1, indicating favorable survival with higher expression, are colored blue. The size of each dot represents the significance level of the survival associations. **e,** Mechanistic model for effect of actin capping protein loss on repair of DNA damage through HDR and vulnerability to CD8 T cell killing.

We observed that for multiple types of cancers there was a significant difference in overall survival (OS) based on the expression of *CAPG*, *CAPZA3* and other HDR associated genes (**Figure 8d**, **Figure 11c-d**). Of the 33 cancer types within the TCGA dataset, 11 showed a significant difference in OS based on *CAPG* or *CAPZA3* expression, as benchmarked with 12 cancer types that showed a significant survival difference based on *BRCA1* or *BRCA2* expression. For most cancers low expression of an HDR associated gene (including *CAPG* and *CAPZA3*) was favorable and was associated with improved OS and an Hazard Ratio (HR) > 1 (**Figure 8d**, **Figure 11c-d**). However, in colorectal cancer (COAD), renal clear cell cancer (KIRC), esophageal cancer (ESCA), stomach cancer (STAD), thyroid cancer (THCA) and thymoma (THYM), the HDR gene had a HR < 1 and low expression was unfavorable and associated with poorer survival (**Figure S11d**). This divergence in patient survival outcomes may reflect the contrasting effects that reduced HDR ability has on cancer cell survival. Low expression of an HDR associated gene may render cancer cells more vulnerable to standard genotoxic therapies such as radiation and chemotherapy; however, it may also enhance cancer cell survival by promoting chronic STING activation and immune suppression (**Figure 8e**). Overall, we found that expression of *CAPG* and *CAPZA3* showed a similar effect on patient OS as other HDR genes; however, the consequence of HDR inhibition will need to be investigated further and will likely differ between cancer types based on tumor specific sensitivity to DNA damage and the mechanisms for immunosuppression.

## Discussion

Through genome-scale CRISPR screens we discovered two actin capping proteins *Capza3*/*CAPZA3* and *CAPG* in cancer cells that mediate differential responses to RT and/or immunotherapy. Validation studies showed that inactivation of these genes induces an HDR defect that leads to persistence of DNA damage and promotes chronic STING activation and the expression of an immune inhibitory ligand, CEACAM1.

Our study identified a previously un-appreciated role of actin capping proteins for efficient HDR in cancer cells. Prior studies have found that DSBs undergoing HDR are very dynamic, clustering into subnuclear compartments that facilitate homology detection and increase HDR efficiency ^23, 34–41^. Formation of these functional domains is believed to involve actin mediated mechanical forces, as nuclear actin forms polymers following exposure to genotoxic agents ^42^ and the expression of a polymerization deficient NLS-R62D actin has been found to reduce HDR rates ^25^. Furthermore, the proteins necessary for actin remodeling in the cytoplasm, Wiskott-Aldrich Syndrome protein (WASP), Arp2/3, formin and actin capping proteins, have also been found within the nucleus ^24,43^ and the reorganization of heterochromatin breaks and efficient HDR is promoted by the formation of actin filaments through Arp2/3 dependent ^39^ and formin dependent mechanisms ^35,42,44^. Actin (β-actin) and actin binding proteins (CAPZβ, APRC4) have also been found to bind to damaged chromatin, and chromatin immunoprecipitation experiments of a specific endonuclease-generated DSB site recovered WASP and ARP2 ^40^, suggesting the interaction of actin-regulating proteins with damaged DNA, either by direct binding or indirectly through other repair factors. Actin and actin binding proteins are not frequently mutated in cancer cells; however, the deregulation of WASP protein, an Arp2/3 activator, leads to HDR deficiency in lymphocytes ^39^ as well as an increased risk for the development of non-Hodgkin’s lymphoma and leukemia ^45^. A recent study also found that RHOJ regulation of formin-dependent nuclear actin polymerization enhanced DNA repair in cancer cells undergoing EMT and promoted their chemoresistance ^46^. These studies, as well as our results here, suggest that profiling and targeting of actin regulating proteins may provide clinically relevant information on HDR deficiency and offer therapeutic insights on the resistance to genotoxic treatments such as chemotherapy and radiation.

We found that *Capza3* KO cells showed increased γH2AX foci and reduced RAD51 foci following radiation exposure. Intriguingly DNA damage specifically within heterochromatin regions showed possible differences in movement between WT and *Capza3* KO cells. In *Capza3* KO cells there was increased γH2AX foci in the nuclear periphery following targeted DNA damage of heterochromatin regions. Movement of DNA DSBs in pericentric heterochromatin to the nuclear periphery prior to HDR is important to avoid aberrant recombination between non-homologous repetitive sequences ^47^. Prior studies in Drosophila have found Rad51 recruitment to DNA DSBs is blocked within the heterochromatin domain and instead Scar and Wash proteins activate Arp2/3 and promote actin polymerization toward the nuclear membrane. These sites of DNA damage are then transported along the actin filament by myosin proteins and anchored to nuclear pores. RAD51 is ultimately recruited to the damaged DNA following actin depolymerization and leads to repair between the damaged DNA and homologous chromosome that were transferred together to the nuclear periphery ^35, 47–50^.

We also found that DNA damage associated with HDR deficiency can activate the STING pathway and lead to immune suppressive downstream effects in addition to the well-known immune enhancing effects. It has consistently been shown that tumors with mutations or epigenetic silencing of genes involved in mismatch repair (dMMR) are vulnerable to ICI ^51–53^. However, ICI use for HDR deficient breast and ovarian tumors has not shown a similar clinical benefit ^54–57^. A possible explanation is that HDR deficiencies may lead to unique immune-suppressive consequences that might impact ICI effectiveness. Activation of cGAS-STING pathway has been previously described in DNA repair deficient cancers following genotoxic stress and shown to induce expression of *PDL1*, a T cell inhibitory ligand ^58–62^. Our study identified that CEACAM1, an inhibitory ligand expressed on cancer cells that can impair T cell function through either homophilic interactions or heterophilic interactions with TIM3, can also be induced following STING pathway activation. In addition to PDL1, our study suggests that CEACAM1 may also be a promising immunotherapeutic target ^63–66^, particularly for HDR deficient cancers, and may lead to more efficacious combination with radiation treatment than has been seen in prior immunotherapy and radiation trials ^67–70^.

## Acknowledgments

We thank the Glazer lab members for reagent sharing and technical assistance. We thank all members of the Chen laboratory, as well as various colleagues at Yale for assistance and/or discussion. We thank the Yale Center for Genome Analysis, High Performance Computing Center, Yale Center for Molecular Discovery, Microscopy Core, and Keck Biotechnology Resource Laboratory at Yale, for technical support.

SC is supported by Cancer Research Institute Lloyd J. Old STAR Award (CRI4964), NIH/NCI (DP2CA238295, R01CA231112, R33CA281702), DoD (W81XWH-20-1-0072, W81XWH-21-1-0514,

HT94252310472), Alliance for Cancer Gene Therapy (ACGT), Pershing Square Sohn Cancer Research Alliance, and YCC Team Science Award. PG is supported by NIH grants (R35CA197574 and R01ES005775). NV is supported by American Board of Radiology’s B. Leonard Holman Research Pathway Fellowship and the ASTRO Radiation Oncology Seed Grant. PAR is supported by Yale PhD training grant from NIH (T32GM007499), Lo Fellowship of Excellence of Stem Cell Research, and YCC T32 fellowship program. CPD is supported by Boehringer Ingelheim Biomedical Data Science Fellowship.

## Author Contributions

Conceptualization: NV, SC. Experiment lead: NV. In-vivo studies: NV, SX. CRISPR Screen and RNA-Seq analysis: PAR. TCGA and patient dataset analysis: CD. Additional experiment support: YF, PR, QL, FZ, NT. Manuscript prep: NV, SX, PG, SC, with inputs from all authors. Supervision: SC, PG. Research funding: SC, NV. Overall organization: NV, SX, PG, SC.

## Methods

### Institutional approval

This study has received institutional regulatory approval. All recombinant DNA and biosafety work were performed under the guidelines of Yale Environment, Health and Safety (EHS) Committee with an approved protocol (Chen-rDNA 21-45). All animal work was performed under the guidelines of Yale University Institutional Animal Care and Use Committee (IACUC) with approved protocols (Chen 20068).

### Mice

Mice of both sexes, between age 8 and 12 weeks, were used for the study. 8-week-old OT-I mice were purchased from the Jackson Laboratory and bred in-house. 8-week-old C57BL/6Nr mice were purchase from Charles River lab. 8-week-old NCr mice were purchase from Charles River lab. All animals were housed in standard, individually ventilated, pathogen-free conditions, with a 12 h:12 h or a 13 h:11 h light cycle, at room temperature (21–23 °C) and 40–60% relative humidity.

### Mouse CD8 T cell culture

Spleens were isolated from OT-I mouse strain and placed in ice-cold wash buffer (2% FBS in PBS). Spleens were dissociated mechanically, passed through a 100 μm filter, and incubated with ACK Lysis Buffer (Lonza) for 2 minutes at room temperature to lyse RBCs. The resulting cell suspension was washed with wash buffer, filtered through a 40 μm filter and then resuspended in MACS Buffer (0.5% BSA and 2 μM EDTA in PBS). Naive CD8^+^ T cells were isolated using naïve mouse CD8 T cell isolation kit from Miltenyi (130-104-075). Naive CD8^+^ T cells were counted and then resuspended in cRPMI (RPMI-1640 with 10% FBS, 2mM L-Glutamine, 1% HEPES, 1% 100 nM NaPyruvate, 1% NEAA, 100U Pen/Strep and .05 µM β-mercaptoethanol) to a final concentration of 1 × 10^6^ cells/ml. The cells were then plated into a 12 well plate at a concentration of 1 × 10^6^ cells per well. These CD8 T cells were expanded *in-vitro* using a previously published protocol ^71^. In brief, cells were expanded for 3 days in cRPMI media supplemented with 10 ng/mL OVA (Anaspec AS-60193-1), 5 ng/mL IL7 and 5 ng/mL IL15. Cytokines were purchased from Peprotech. After 3 days of *in-vitro* expansion CD8 T cells were used for co-culture experiments.

### Generation of B16F10 cells that express Cas9 and OVA antigen

We generated a B16F10 cell line that expresses Cas9 protein and OVA antigen to be used in our screen. For all experiments B16F10 cells were maintained in complete DMEM media (DMEM with 10% FBS, 2mM L-Glutamine and 100U Pen/Strep). Cells were passaged when they reached confluence, which was typically 2-3 days after splitting depending on the splitting ratio. Cells were passaged using TrypLE and cell lines were frozen using DMSO-containing media.

To generate a Cas9 expressing B16F10 line (B16F10-Cas9) a lentiviral Cas9-Blast vector (Addgene Plasmid # 52962) was co-transfected with packaging plasmids PAX2 and pMD2.G into HEK293T. Transfection was performed using LipoD293T (Signagen # SL100668) per the manufacturer’s protocol. Virus was harvested at 48 hours post-transfection, tittered, and stored at −80°C. B16F10 cells were transduced with Cas9-Blast lentivirus overnight. Two days later, infected cells were selected with 5 µg/ml of blasticidin for at least 3 days. Clonal lines of B16F10-Cas9 were established and their cutting efficiency was verified using *Pdl1* targeting gRNA and flow cytometry analysis of PD-L1.

Next, to overexpress OVA antigen within the B16F10-Cas9 line, lentivirus that contain an OVA-mCherry vector (Sidi Chen lab) were generated using the procedure described above. The B16F10-Cas9 clonal lines were then transduced with this lentivirus and 2 days after transduction mCherry positive cells were sorted, expanded and frozen down. OVA expression was verified using OVA-MHCII (Biolegend 141605) flow analysis. The B16F10-Cas9-OVA cell line was confirmed negative for mycoplasma by quantitative RT-PCR.

### Human cell culture

MDA-MB-231 (ATCC HTB-26) and PANC1 (ATCC CRL-1469) cell lines were cultured in complete DMEM media (DMEM with 10% FBS, 2mM L-Glutamine, and 100U Pen/Strep). Cells were passaged when they reached confluence, typically 2-3 days after splitting, depending on the splitting ratio. Cells were passaged using TrypLE and cell lines were frozen using DMSO-containing media. All human cell lines were confirmed negative for mycoplasma by quantitative RT-PCR.

### B16F10 cell culture and chemical Treatment with Arp2/3 inhibitor (CK-666) and STING inhibitor (H151)

B16F10 (ATCC CRL-6475) cell lines were cultured in complete DMEM media (DMEM with 10% FBS, 2mM L-Glutamine, and 100U Pen/Strep). Cells were passaged when they reached confluence, typically 2-3 days after splitting, depending on the splitting ratio. Cells were passaged using TrypLE and cell lines were frozen using DMSO-containing media. All cell lines were confirmed negative for mycoplasma by quantitative RT-PCR. For experiments using Arp2/3 inhibitor CK-666 (Fisher) and STING inhibitor H151 (Abcam), the inhibitors were added to B16F10 cells 24 hours prior to radiation at varying concentrations (10 μM, 50 μM and 100 μM).

### Genome scale CRIPSR screen in B16F10 cells

#### gRNA pool library production

Mouse CRISPR Brie lentiviral pooled library (Addgene Plasmid # 170511) consisting of 79,637 gRNAs was co-transfected with packaging plasmids (pPAX2 and pMD2.G) into HEK293T cells using LipoD 293T transfection reagent (Signagen # SL100668) following the manufacture’s protocol. 24 hours after transfection the media was replaced. Virus supernatant was collected at 48 and 72 hours after transfection. Virus was concentrated using PEG virus precipitation (Promega # V3011). In brief, the collected supernatant was pooled and spun down at 3000g for 15 minutes to remove cell debris. Supernatant was carefully collected and 8 mL of 40% PEG8000 solution was added to 32 mL of viral supernatant for a final concentration of 8% PEG8000. After mixing well with vortexing the virus supernatant was incubated overnight at 4C. The next day the viral supernatant was centrifuged at 3000g for 30 minutes at 4C. The supernatant was aspirated carefully to avoid disturbing the virus pellet which was then resuspended in cRPMI, divided into small aliquots and stored at −80°C. To determine virus titer 1×10^6^ B16F10-Cas9 cells were plated per well of a 6-well plate. B16F10 cells were transduced with different amounts of the aliquoted lentivirus in the presence of 8 µg/ml of polybrene. The next day, transduced B16F10 cells from each condition were seeded at a density of 10,000 to 100,000 cells per well of a 6-well plate (in triplicates). Twenty-four hours following infection, puromycin (2µg/ml) was added. After 3 days of puromycin selection infected cells in each well were counted.

#### Screen

B16F10-Cas9-OVA cells (hereafter referred to as B16F10) were transduced with the Brie library lentivirus at a MOI of 0.3 and cells were selected with puromycin (2 µg/mL) for 3 days prior to use in co-culture with CD8 T cells. Mouse CD8 T cell isolation was performed as described above. Prior to co-culture approximately 120 million B16F10 cells were isolated as a baseline control. There were four treatment conditions tested in our screen: 1) No treatment, 2) anti-PD1 antibody during co-culture, 3) 1 Gy radiation before co-culture, and 4) 1 Gy radiation before co-culture followed by co-culture with anti-PD1 antibody. To maintain at least a 1000x fold coverage, approximately 100 million B16F10 cells were used for each condition. B16F10 cells were plated into 24 well plates (1 million cells per well) and pretreated with 10 ng/ml of IFN-γ for 24 hours prior to co-culture with OT-I CD8 T cells to increase MHC class II expression. For treatment conditions that included radiation the plated B16F10 cells were exposed to 1 Gy radiation (using MultiRad350 irradiator per the manufacturer’s protocol) 1 hour before the addition of OT-I CD8 T cells. For conditions that included anti-PD1 antibody treatment we used anti-mouse PD-1 inVivo mAB # BE0146 (clone: RMPI-14, Lot#806321A2B) at a concentration of 10 µg/mL. With the addition of CD8 T cells to B16F10 the media was changed to cRPMI with IL7 (5 ng/mL) and IL15 (5 ng/mL). Co-culture was done at a 1:1 T:E ratio. 1 day after co-culture we performed Annexin V bead purification (Miltenyi # 130-090-201) per the manufacturer’s protocol. Cells were passed through the column twice to increase purification of Annexin V negative cells. 2 replicates of the screen, starting with lentiviral infection of B16F10 cells with the Brie library, were done.

#### Library preparation and sequencing

Genomic DNA was isolated from Annexin V negative cells using the DNeasy Blood and Tissue kit (Qiagen # 51192) following the manufacturer’s protocol. PCR amplification of the gRNA cassette for Illumina sequencing of gRNA representation was done using the Broad protocol available online (https://media.addgene.org/cms/filer_public/56/71/5671c68a-1463-4ec8-9db5-761fae99265d/broadgpp-pdna-library-amplification.pdf). NGS Illumina sequencing was done by the YCGA core to a depth of 200x.

#### Screen data analysis

Raw sequencing fastq data had adapter sequences trimmed via Cutadapt v3.41 ^72^ using a 10% error rate and the following sequences: forward, 5’-tcttgtggaaaggacgaaacaccg; reverse, 5’-gttttagagctagaaatagcaagt. Trimmed sequences were then filtered to remove those with <15 nt length. The remaining sequences were aligned to a reference, comprised of the CRISPR sgRNA-spacer sequences. Alignment was performed using Bowtie v1.3.02 with the following settings: -v0, -m1 –best. The sgRNA counts for each sample were processed and analyzed using SAMBA R package v1.3.0 (https://github.com/Prenauer/SAMBA) (detailed below) ^73^. Specifically, sgRNAs were filtered to include those with >10 counts across screened samples (non-control). A two-step data analysis was performed, first with an sgRNA-level analysis by the edgeR R package v3.38.43 pipeline with TMM-wsp size factors, feature-wise dispersion, quasilikelihood (QL) generalized linear model fitting, and QL F tests. In the second analysis step, sgRNA scores were aggregated into a gene score, calculated as a weighted sum of the sgRNA log2 fold-changes (log2-FC). Gene level p values were assessed based on a null distribution of gene scores, which were scored from randomly grouped sgRNAs of non-targeting controls. P values were adjusted using the method by Benjamini and Hochberg. An additional metric to assess gene enrichment was the number of sgRNA / gene with a log2-FC > the 90th percentile of the randomized null data log2-FC, representing a 10% FDR. Screen data was also analyzed with the commonly used MAGeCK RRA algorithm for robust comparison ^20^.

The screen analyses included all samples in a statistical model that incorporated cocultured B16F10 that were treated with anti-PD-1 and / or radiation therapy (RT) (∼ Coculture + anti-PD-1 + RT + anti-PD-1&RT). From this statistical model, we separately assessed the effects of anti-PD-1, RT, and the combined treatment coefficients with SAMBA. We verified that there was high gRNA detection for all treatment conditions in both screen replicates and that the treatment conditions for the two replicates showed high correlation and clustered together on PCA component analysis.

### Generation of B16F10 *Capza3* KO cell lines

To generate the B16F10 *Capza3* KO lines the 2 CRISPR gRNAs enriched in the radiation alone and radiation and anti-PD1 antibody screen treatment conditions were cloned into the LRG vector (Addgene Plasmid # 65656). For WT controls a NTC gRNA from the CRISPR screen was cloned into the LRG vector. Lentivirus was made by co-transfecting these vectors with packaging plasmids (PAX2 and pMD2.G) into HEK293T cells using LipoD 293T transfection reagent (Signagen # SL100668) following the manufacturer’s protocol. B16F10 cells were transduced with lentivirus using the protocols described above. 2 days after transduction, GFP positive B16F10 cells were sorted into a 96 well plate at a density of 10 cells per well and allowed to expand. Each well thus contained a mixture of clonal lines. To enrich B16F10 cells that had mutations at the *Capza3* loci we sequenced each of the wells to identify those that had depleted WT sequences by sanger sequencing. These wells were expanded and then the cells sorted again (into a 96 well plate at a density of 10 cells per well) with a second round of sequencing to identify wells which had absence or a low proportion of WT cells. Using this process, we generated 2 mixtures of clonal KO cells for *Capza3* gRNA 2 targeting (*Capza3* gRNA 2 #1 and *Capza3* gRNA 2 #2) and 2 mixtures of clonal KO cells using *Capza3* gRNA 3 targeting (*Capza3* gRNA 3 #1 and *Capza3* gRNA 3 #2). To maintain the same passage number, WT cells transduced with LRG containing a NTC gRNA went through the same procedure described above. Nextera library preparation and next generation sequencing was used to quantify the percent modified alleles in both Capza3 gRNA targeting loci in our KO mixtures and in WT cells and are shown in a supplemental figure. Oligonucleotide sequences are listed in Supplementary Tables.

### Nextera library preparation and sequencing

The region around both *Capza3* gRNA targeting loci were amplified by PCR and then tagged, amplified, and barcoded using Nextera XT DNA Library Prep Kit (Illumina) per the manufacturer’s protocol. The library for each sample was quality controlled and quantified separately using the 4150 TapeStation System (Agilent), followed by library pooling and PCR clean up using QIAquick PCR Purification Kit (Qiagen). Libraries were denatured and diluted to 10 pM according before loading on MiSeq (Illumina) for next generation sequencing. FASTQ reads were quality controlled by running FastQC v0.11.9 ^74^ and contaminations by Nextera transposase sequence at 3’ end of reads were trimmed using Cutadapt v3.2 ^72^. Processed reads were aligned to amplicon sequence and quantified for insertions, deletions (indel), and substitution using CRISPResso2 v2.1.3 ^75^. Specifically, we retrieved amplicon sequences, which were 150 ∼ 250 bp (according to the length of reads) flanking crRNA target sites, from the mm10 genome. A 5-bp window, centered by predicted Cas9 cutting sites, was used to quantify genetic modification for each crRNA in both WT and *Capza3* KO cells. Percent modification and allele frequency plots were generated with CRISPResso2 v2.1.3.

### Immunofluorescence of DNA damage markers and quantification

High-throughput immunofluorescence foci assays were performed at the Yale Center for Molecular Discovery (YCMD). Polystyrene flat bottom 384-well plates (CellVis # P384-1.5H-N) were coated with collagen O/N at 4C (10 ng/mL diluted in PBS). Collagen was aspirated and cells (B16F10, PANC1, MDA-MB-231, SKMEL28) were seeded at 10,000-50,000 cells/well and allowed to adhere overnight. Cells were irradiated (using MultiRad350 per manufacturer’s protocol) and then fixed at different time points (1 hour, 3 hours, 6 hours and 24 hours) before staining for DNA damage markers: γH2AX (Cell Signaling Technology # 9718T), 53BP1 ((Novus Biologicals # NB100-904) and phospho-RPA2 (pRPA, Bethyl Laboratories # A300-246A). Cells were fixed with 4% paraformaldehyde in PBS for 15 min, washed twice with PBS, and then incubated in permeabilization buffer (0.3% Triton X-100 in PBS) for 15 min. Cells were washed twice with PBS and then incubated in blocking buffer (2% BSA in PBS) or 1 h. All primary antibodies were incubated O/N at 4°C at the appropriate dilution: γH2AX (1:200), 53BP1 (1:200), pRPA (1:250). After primary antibody staining the cells were washed three times with PBS and then incubated with Alexa Fluor dye conjugated secondary antibody (ThermoFisher # A-11012) and DAPI (Biolegend # 422801) for 1 hr at RT. After secondary antibody staining cells were washed 3 times with PBS and images were taken on either the InCell Analyzer 2200 Imaging System or the Leica SP8 confocal microscope. Quantification of DNA damage foci was performed by YCMD using the Incell Analyzer software.

Additional small-scale immunofluorescence experiments were performed using 8-well chamber slides (Ibidi # NC0704855). These slides were coated with collage O/N at 4C (10 ng/mL diluted in PBS). Collagen was aspirated and cells (B16F10, PANC1, MDA-MB-231, SKMEL28, A375) were seeded at 100K-200K cells/well and allowed to adhere overnight. Following irradiation (using MultiRad350 per manufacturer’s protocol), cells were fixed and stained for γH2AX, 53BP1, RAD51 and pRPA as described above. Images were taken using the Leica SP8 confocal microscope at 63X magnification.

### Colony forming assay (CFA)

B16F10 cells were plated 1 day before they were irradiated at varying doses of ionizing radiation (using MultiRad350 per manufacturer’s protocol). Four to six hours after irradiation, cells were detached using TrypLE, washed with complete DMEM media, counted, and seeded in 6-well plates in triplicate at a density of 100 cells per well. These plates were kept in the incubator for 10 to 14 days. After incubation, colonies were washed in PBS, fixed with ice-cold 100% methanol for 10 minutes at −20C and then stained with crystal violet (0.5% crystal violet in 25% methanol). Colonies were counted using a brightfield microscope. Colonies needed to have at least 10 cells to be counted as a colony. For PANC1 and MDA-MB-231 cells lines irradiation was carried out in the same way as described above. However, to make sure the cell number was accurate we used FACS sorting to sort 50 cells per well into a 24 well dish. Staining with crystal violet and quantification was carried out as described above.

For colony forming assay after drug incubation, cells were cultured for 1 day with different concentrations of the drugs cisplatin (Selleckchem #S1166), AZD7648 (DNA PK inhibitor, Cayman # 28598) and Olaparib (Parp inhibitor, Cayman # 10621-25). 1 day after culture the cells were detached using TrypLE, washed with wash buffer (2% FBS in PBS) and then FACS sorted 50 cells per well into a 24 well dish. Staining with crystal violet and quantification was carried out as above.

### Extra-chromosomal luciferase reporter assays

The NHEJ and HDR luciferase reporters has been previously reported ^26^ and were obtained from the Peter Glazer lab. For the HDR luciferase reporter, a DSB in the firefly luciferase gene was induced by I-SceI digestion and confirmed by electrophoresis. Linearized plasmid was then transfected into cells. HDR within the cells will restore the luciferase activity, which can then be used to measure HDR efficiency. To assay NHEJ, a HindIII-mediated DSB was generated within the NHEJ luciferase reporter and confirmed by electrophoresis. After transfection of the linearized NHEJ plasmid, repair of the DSB by NHEJ restores firefly luciferase activity.

All reporter assays were performed in a 96-well format by seeding 50,000-100,000 cells per well, 24 hours before transfection. Lipofectamine 3000 (ThermoFisher # L3000008) was used for transfection per the manufacturer’s protocol. As a control for transfection rates a plasmid with an intact luciferase was used for each cell line in a separate well. Luciferase following transfection with linearized HDR and NHEJ reporters was then normalized to this control for each sample. Luciferase activity was read using a bioluminescence plate reader.

### Laser Micro-irradiation

Laser micro-irradiation experiments were done using 8-well chamber slides (Ibidi # NC0704855). These slides were coated with collage O/N at 4C (10 ng/mL diluted in PBS). Collagen was aspirated and B16F10 were seeded at 100K-200K cells/well and allowed to adhere overnight. Cells were incubated with 1 µg/mL Hoechst nucleic acid dye for 2 hours before the imaging experiment. Cells were mounted on a Leica TCS SP8 X microscope system (Leica Microsystems) with an incubator chamber at 37°C with 5% CO_2_. For laser micro-irradiation cells were exposed to a 405 nm diode laser (95% with FRAP booster, 35 iterations, 150 µJ/pixel total power) and irradiation field of multiple 5-pixel wide stripes. For experiments in which different nuclear regions were targeted we targeted Hoechst bright areas as a proxy for heterochromatin, the nucleolus (Hoechst dim) as a proxy for euchromatin and the nuclear membrane as a control. At different time points after irradiation (1 minute, 5 minutes, 10 minutes, 15 minutes, 20 minutes, and 2 hours), cells were fixed with 4% PFA in PBS for 10 min at RT. γH2AX staining was done as described above. For analysis of micro-irradiation stripe intensity ImageJ was used.

### Generation of B16F10 H2B mCherry cell lines and live cell imaging

To generate B16F10 cells with a H2B-mCherry reporter the H2B mCherry reporter plasmid (Addgene Plasmid # 20972) was co-transfected with packaging plasmids PAX2 and pMD2.G into HEK293T. Transfection was performed using LipoD293T per the manufacturer’s protocol. Virus was harvested at 48 hours post-transfection and stored at −80°C. WT and *Capza3* KO B16F10 cells were transduced with lentivirus and two days later mCherry positive cells were sorted and expanded.

Live cell imaging was performed using 8-well chamber slides (Ibidi # NC0704855). These slides were coated with collagen O/N at 4C (10 ng/mL diluted in PBS). Collagen was aspirated and B16F10 were seeded at 100K-200K cells/well and allowed to adhere overnight. Cells were mounted on a Leica TCS SP8 X microscope system (Leica Microsystems) with an incubator chamber at 37°C with 5% CO_2_. In addition to the H2B-mCherry reporter we also used Hoechst nucleic acid dye (ThermoFisher # H3570) to visualize micronuclei formation. Cells were incubated with 1 µg/mL Hoechst nucleic acid dye for 2 hours before the imaging experiment. Media was replaced with normal culture media prior to imaging. For the imaging, 10 fields of view were taken for each sample over a span of 24 hours. To quantify micronuclei formation, we used the Leica SP8 LasX software. We used both Hoechst and H2B mCherry to identify the nucleus and determine if there were any discrete DNA aggregates separate from the nucleus that may represent micronuclei. Identified micronuclei showed both Hoechst staining and H2B-mCherry signal. Cells with nuclei associated with at least 1 micronucleus were considered positive.

#### Generation of MDA-MB-231 and PANC1 *CAPG* and *CAPZA3* KO cell lines

*CAPG* and *CAPZA3* targeting gRNAs to generate the human KO lines were identified using CRISPick (https://portals.broadinstitute.org/gppx/crispick/public). These gRNAs were cloned into a plasmid containing Cas9 and a BFP reporter (Addgene Plasmid # 64216) and cutting efficiency was tested in 293T after transient transfection using lipoD293T and the T7E1 assay. The gRNA with the highest cutting efficiency was used for the targeting experiments. In brief, MDA-MB-231 and PANC1 cell lines were dissociated into single cells and replated at a density of 1 million cells per well into a 12 well plate. The cells were then transfected with the BFP reporter plasmid containing *CAPG* and *CAPZA3* targeting gRNAs using Lipofectamine 3000 (ThermoFisher # L3000008). 2 days after transfection the cells were sorted and plated as individual cells into each well of 96 well plates. Cells were expanded and then divided into 1 plate for continued culture and 1 plate for colony PCR and sequencing. DNA Quick Extract (Lucigen # 76081-766) was used to isolate DNA. The *CAPG* and *CAPZA3* loci were then amplified by PCR and submitted for Sanger sequencing. Those clones that showed mutations at these loci were expanded and frozen down. For all targeting experiments we also isolated and froze down lines that had undergone the targeting procedure, but whose genomic sequence at *CAPG* and *CAPZA3* was not changed, these were used as our passage-matched WT controls. Oligonucleotide sequences are listed in Supplementary Tables.

#### Generation of *CEACAM1* KO cell lines

*CEACAM1* targeting gRNAs to generate the mouse and human KO lines were identified using CRISPick (https://portals.broadinstitute.org/gppx/crispick/public). For B16F10 cells, a gRNA targeting *Ceacam1* was cloned into a lentiviral vector containing the puromycin selection marker. Lentivirus was generated as described above and cells were transduced and selected with puromycin for 5 days. Targeting of *Ceacam1* was confirmed by performing flow analysis of Ceacam1 prior to performing experiments. To generate the human KO lines gRNAs were cloned into a plasmid containing Cas9 and a BFP reporter (Addgene Plasmid # 64216) and cutting efficiency was tested in 293T after transient transfection using lipoD293T and the T7E1 assay. The gRNA with the highest cutting efficiency was used for the targeting experiments. PANC1 WT, *CAPG* KO and *CAPZA3* KO cell lines were used for targeting. Transfection, sorting, expansion, and sequencing were carried out as described above to generate single *CEACAM1* KO, *CEACAM1*/*CAPG* DKO and *CEACAM1*/*CAPZA3* DKO cell lines. We also isolated and froze down WT, *CAPG* KO and *CAPZA3* KO cell lines that had undergone the *CEACAM1* targeting procedure, but whose genomic sequence at *CEACAM1* was not changed, these were used as our passage-matched controls.

### Generation of PANC1 *AAVS1* TLR cell line and *CAPG*, *CAPZA3* KOs, and testing HDR/NHEJ efficiency

Traffic light reporter (TLR) constructs were ordered from Addgene (Plasmid #s 64323, 64322, 64216, and 64215) and transfected into PANC1 using Lipofectamine 3000 (ThermoFisher # L3000008) per the manufacturer’s protocol. We decided to generate the TLR reporter in the PANC1 background as PANC1 cells showed higher transfection efficiencies than MDA-MB-231. 1 day following transfection of a plasmid containing a BFP reporter and the *AAVS1* targeting gRNA and the pAAVS1-TLR targeting plasmid we sorted BFP positive cells into 1 cell per well of a 96 well plate. After the cells had expanded, genomic DNA was isolated for colony PCR. After identifying and expanding the targeted clones we performed allele specific PCR and found that both of our targeted PANC1 lines had homozygous insertions of the TLR construct into the *AAVS1* locus. We verified that after transfection with the pU6-sgRosa26-1_CBh-Cas9-T2A-BFP plasmid (Addgene Plasmid # 64216) which contains a *Rosa26* targeting gRNA, the cells expressed RFP and GFP. These lines (PANC1 *AAVS1* TLR) were expanded and frozen down. *CAPG* and *CAPZA3* KOs were generated in these lines using the protocols described above. As described above, passage-matched WT controls were those PANC1 *AAVS1* TLR cells that went through the same transfection with *CAPG* and *CAPZA3* gRNAs and clonal expansion procedure but did not show any mutations at the *CAPG* or *CAPZA3* locus.

To analyze HDR and NHEJ efficiency using the TLR reporter, we transfected the pU6-sgRosa26-1_CBh-Cas9-T2A-BFP plasmid (Addgene Plasmid # 64216) which contains a *Rosa26* targeting gRNA and the pTLR repair vector (Addgene Plasmid # 64322) into WT, *CAPG* KO and *CAPZA3* KO PANC1 *AAVS1* TLR lines. Transfection was done using Lipofectamine 3000 (ThermoFisher # L3000008), based on the manufacturer’s protocol. 1 day after transfection the media was replaced. We performed flow analysis by gating on BFP positive cells (which represented transfected cells) and then quantifying the precent GFP and RFP positive cells at 24, 48 and 72 hours after transfection.

### RNA-seq

For RNA-seq, total RNA was isolated with the RNeasy Mini Kit (Qiagen # 74136) from B16F10 WT and *Capza3* KO cells, before and after radiation treatment (1 Gy). The *Capza3* KO cell lines were generated as described above. RNA was collected 1 day after radiation treatment. We performed transcriptome profiling using RNA-seq in WT and *Capza3* KO B16F10 cells, with 2 independent clones from each genotype/group (WT, *Capza3* KO gRNA 2, and *Capza3* KO gRNA 3), with biological duplicates for each clone (2 x 2 = 4, quadruplicates for each genotype). The cells were analyzed before and after radiation treatment, for a total of 24 samples. RNA samples were submitted to the YCGA core for library prep and sequencing.

Sequencing data were aligned to the mouse genome (GRCm39) using the STAR aligner v2.7.11 ^76^. Briefly, an alignment reference panel was created from the Gencode vM32 primary genome assembly with a sjdbOverhang of 149 (read length – 1). Alignment was performed using the “TranscriptomeSAM” quantification mode from STAR, and transcript counts were estimated using RSEM v1.3.3 ^77^ with the “rsem-calculate-expression” function. The gene count data were filtered and analyzed using the edgeR R package pipeline ^78^ with a 2-step upper quartile normalization ^79^, and the full statistical model included the following covariates: gene-targeting guides (GT) and gRNA, given that two different gRNAs were used (∼ GT + gRNA). A likelihood ratio test was used to compare B16F10 cells treated with gene-targeting and non-targeting gRNAs. Transcription factor activity prediction was performed using the decoupleR R package v2.2.2 ^80^ with the run_fgsea function, the DoRothEa database ^81^ for directional interactions between TF/target-genes from the (mouse data; A-C interaction confidence), and 1000 iterations.

### Quantitative-PCR

Total RNA was isolated with the RNeasy Mini Kit (Qiagen # 74136). DNA was removed from RNA samples using genomic DNA eliminator spin columns. cDNA was produced from RNA using SuperScript III Reverse Transcriptase kit (Life Technologies # 18080051) or High-Capacity cDNA Reverse Transcriptase kit (Life Technologies # 4368813). Quantitative real-time PCR was performed in triplicate using PowerTrack QPCR SYBR Green (ThermoFisher # A46109). Oligonucleotide sequences are listed in a Supplementary Table.

### Flow cytometry analysis

#### Surface marker

Cells were disaggregated with TrypLE for 5 minutes and washed with cold wash buffer (2% FBS in PBS). Cells were pelleted by centrifugation and washed again with wash buffer. Each sample was resuspended in wash buffer with the appropriate conjugated antibody and LD APC-Cy7 (Thermo # L34976). Cells were incubated in wash buffer with the antibody for 30 minutes on ice. After staining cells were washed two times with wash buffer and resuspended in wash buffer and analyzed by FACS. A complete listing of antibodies used is presented in the key reagents section below. Representative flow plots are shown where appropriate.

#### Annexin V staining

Supernatant was collected and cells were detached following incubation with 1 mM EDTA for 5 min at RT. Cells were washed twice with wash buffer by centrifuging at 300g for 10 minutes. Cells were then resuspended in 90 µL of 1x Binding Buffer per 10^7^ cells and 10 µL of Annexin V-APC antibody and DAPI was added. Cells were incubated for 15 minutes at 4C before washing twice with 1x Binding Buffer and centrifugation at 300g for 10 minutes. Cells were then analyzed by FACS.

#### Intracellular staining (phosphor-TBK1, phosphoIRF3, STING)

24 hours after radiation treatment cells were disaggregated with TrypLE for 5 minutes and washed with cold wash buffer (2% FBS in PBS). Cells were pelleted by centrifugation and washed again with wash buffer. Each sample was resuspended in wash buffer with LD APC-Cy7 and incubated for 30 minutes on ice. After staining cells were washed two times with wash buffer and then fixed with BD Cytofix Buffer (BD Biosciences # 554714) for 20 minutes on ice. Following fixation cells were washed two times with wash buffer before permeabilization with ice cold 100% methanol for 1 hour on ice. After permeabilization cells were washed two times with wash buffer (after permeabilization cells were centrifuged at 800xg to avoid cell loss), before staining with phosphor-TBK1 (Cell Signaling # 13498S), phosphor-IRF3 (Cell Signaling # 83611S) and STING (Cell Signaling # 78827S) for 1 hour on ice. After staining cells were washed two times with wash buffer and resuspended in wash buffer and analyzed by FACS.

### EGFR-CAR-T cell generation

Cryopreserved PBMCs were obtained from StemCell Technologies (Catalog # 70025). PBMCs were thawed and CD3+ T cells were isolated immediately by Pan T Cell Isolation Kit (Miltenyi # 130-096-535). Then isolated T cells were incubated by CD3/CD28 Dynabeads (Thermofisher # 11131D) for T Cell expansion and activation for 48 hours. Following activation, the T cells were transduced with lentiviral generated using EFS-scFv-CD8TM-41BBL-CD3zeta-T2A-Puro-WPRE (Sidi Chen lab). The T cells were transduced by spin-infection, in which 50 µL of concentrated lentivirus was added to 1 million activated CD3+ T cells in 1 ml of complete media (X-VIVO with 5% AB Serum and IL2) with 8 µg/ml of polybrene.

Cells with lentivirus were spun at 900g at 37C for 90 minutes, followed by incubation for 24 hours in 37C incubator. Following incubation, virus was removed and puromycin (1ug/ml) was add for 5 days to get successfully transduced CAR-T cells. Expression of the CAR receptor was confirmed using Flag flow cytometry.

### PANC1/MDA-MB-231 Co-culture

PANC1 and MDA-MB-231 cells were plated in 48 well plates for 24 hours prior to the addition of EGFR CAR-T cells. PANC1 cells were plated at a density of 100K cells/well and MDA-MB-231 were plated at a density of 50K cells/well. 6 hours after plating the cells were irradiated and 24 hours after irradiation we added the EGFR CAR-T at a 1:1 T:E ratio. We observed significant cell death by 12 hours for the PANC1 co-culture and by 24 hours for the MDA-MB-231 co-culture. Thus, these were the time points we decided to halt the experiment and quantify the number of surviving cancer cells using FACS. In brief, all cells (attached and detached) were collected. The cells were washed 2x in cold wash buffer (2% FBS in PBS). Cells were stained with LD APC-Cy7 (Thermo # L34976) and CD3 FITC (Biolegend # 300406). For all experiments a sample of cancer cells alone and EGFR Car-T cells alone were used as gating controls. Using FACS we counted the total number of live cells within a sample and then analyzed the number of these live cells that were CD3 negative to obtain the number of surviving cancer cells. Representative FACs plots showing gating and CD3 staining are show in a supplemental figure. For some experiments anti-human TIM3 antibody (Biolegend # 345009, 10 μg/ml) or anti-human PD1 antibody (BioXCell # BE0188, 10 μg/ml) was added during co-culture with EGFR CAR-T for blocking/neutralization.

### B16F10 *in-vitro* co-culture with OT-I CD8 T cells

For the OT1 T cell killing assay B16F10-Ova cells were cultured with OT1 CD8 T cells. OT1 CD8 T cells were isolated following the instructions of the Naive CD8 T Cell Isolation Kit (Miltenyi Biotec) and activated as described above. OT1 CD8 T cells were added to B16F10 cells 8 hours after the B16F10 cells were plated. The tumor cells and OT1 T cells were co-cultured at tumor: effector ratios of 1:1 and 1:2. Tumor cell killing was tested 24 hours after addition of OT1 CD8a^+^ T cells using GFP or as a time course up to 72 hours after plating B16F10 cells using xCELLigence RTCA eSight live cell analysis platform.

### Testing of B16F10 WT and *Capza3* KO cell lines in syngeneic tumor models

For the B16F10 melanoma model, 1×10^^6^ B16F10 cancer cells were subcutaneously injected into the right flank of nCR mice and C57BL/6 mice. The tumor volume was measured every 2 days. 8 days (nCR) or 17 days (C57BL/6) post transplantation the subcutaneous tumors in the right flank were irradiated with 5-10 Gy radiation with a custom lead shielding used to protect the rest of the mice from radiation. Tumor volume was calculated with the formula: volume = π/6*xyz. Two-way ANOVA was used to compare growth curves between treatment groups. 24 hours post radiation (nCR) or 24 days post transplantation (C57BL/6) tumors were harvested for analysis of STING pathway activation and CEACAM1 expression (nCR tumors) and tumor infiltrated immune cells (C57BL/6 tumors).

### Analysis of tumor-infiltrated immune cells

Tumors were harvested 24 days post transplantation and minced into 1 mm size pieces and then digested with 100U/ml collagenase IV and DNase I for 60min at 37°C. Tumor suspensions were filtered through 100-μm cell strainer to remove large bulk masses. The cells were washed twice with wash buffer (PBS plus 2% FBS). 1 ml ACK lysis buffer was added to lysis red blood cell by incubating 2-5 min at room temperature. The suspension was then diluted with wash buffer and spun down at 400g for 5min at 4°C. Cell pellet was resuspended with wash buffer and followed by passing through a 40μm cell strainer. Cells were spun down and washed twice with wash buffer. Finally, cell pellets were resuspended in MACS buffer (PBS with 0.5% BSA and 2mM EDTA). The single cell suspensions were used for flow cytometry staining and FACS sorting. For immune cell staining first CD45^+^ cells were gated, then particular immune cell types were quantified: CD8 T cells (CD45^+^, CD3^+^, CD8^+^), CD4 T cells (CD45^+^, CD3^+^, CD4^+^), and macrophages (CD45^+^, CD11b^+^, F4/80^+^). The percentage of LAG3 positive T cells (CD45^+^, CD3^+^, LAG3^+^) and PD1 positive T cells (CD45^+^, CD3^+^, PD1^+^) were also quantified. All flow antibodies were used at 1:200 dilution for staining. The LIVE/DEAD Near-IR was diluted 1:1000 to distinguish live or dead cells. Gating for flow analysis is shown in (**Figure S10b**)

### TCGA analysis

We retrieved processed mRNA expression profiles from the TCGA GDC data portal (https://portal.gdc.cancer.gov/) using TCGAbiolinks with the following options: data_category: Transcriptome Profiling, data_type: Gene Expression Quantification, experimental_strategy: RNA-Seq, workflow_type: STAR-Counts, data_format: TPM_unstrand. The curated clinical and somatic mutation information (MuTect2 Variant Aggregation and Masking pipeline) was downloaded from UCSC Xena ^82^. Tumor mutation burden was defined as the count of nonsynonymous mutations, including missense, nonsense, frameshift, and splice-site mutations. Proportions of tumor-infiltrating immune cells were estimated from gene expression data using the ‘MCPcounter’ package. Differentially expressed genes between high and low CAPG/CAPZA3 expression patients were identified using the ‘limma’ package. Pathway enrichments were conducted in R (v4.3.0) using the ‘clusterProfiler’ package ^83^. For analysis of the effect of CAPG expression on ICI efficacy, Van Allen et al. 2015 ^84^ dataset with 121 melanoma patients receiving ICI was used. For analysis of CART response dataset GSE100797 was used. For analysis of radiation sensitivity the following datasets were used: GSE168009, GSE186940 and GSE226034.

### Statistical Analysis

Data are presented as means ± SD, unless otherwise noted. Data was compared using Student’s *t* test, or ANOVA with repeated measures when appropriate. The test used is indicated in the figure legends. All tests were two-sided. Statistical analyses were carried out using GraphPad Prism. A p value of less than 0.05 was considered statistically significant.

### Illustrations

Illustration of schematics were performed using Affinity Designer and Biorender (https://www.biorender.com).

### Reporting summaries Statistics

For all statistical analyses, we confirmed that the items mentioned in NPG reporting summary are present in the figure legend, table legend, main text, or Methods section.

### Software and code

#### Data collection

Flow cytometry data were collected by Attune NxT Flow Cytometer (Thermo), Four-laser Aria II (BD), Five-laser Symphony S6 (BD), Cytek Aurora (Cytek Biosciences); All the deep sequencing data were collected by Yale Center for Genome Analysis (YCGA).

### Data analysis

Data analysis was performed using the following software / packages:

Prism 10.1.0; FlowJo v.10.9.0; Prism 9; CRISPResso2 v2.1.3; Bowtie 1.3.0; Cutadapt v3.4.0; EdgeR v3.38.4; stringr v1.5.0; dplyr v1.1.1; ggplot2 v3.4.1; ggrastr v1.0.1; ggrepel v0.9.3; patchwork v1.1.2; cowplot v1.1.1; reshape2 v1.4.4; factoextra v1.0.7; limma v3.52.4; cluster v2.1.4; DESeq2 v1.36.0; SAMBA v2.0

### Standard statistical analysis

All statistical methods are described in figure legends and/or supplementary Excel tables. The *p* values and statistical significance were estimated for most analyses. One-way ANOVA, two-way ANOVA, Dunnett’s multiple comparisons test, Tukey’s multiple comparisons test was used to compare multiple groups. Data between two groups were analyzed using a two-tailed unpaired *t*-test. Different levels of statistical significance were accessed based on specific *p* values and type I error cutoffs (0.05, 0.01, 0.001, 0.0001). Data analysis was performed using GraphPad Prism v.10. and RStudio. Source data and statistics were provided in a supplemental excel table.

### Data and resource availability

All data generated or analyzed during this study are included in this article and its supplementary information files. Specifically, source data and statistics for non-high-throughput experiments such as flow cytometry, protein experiments, and other molecular or cellular assays are provided in an excel file of Source data and statistics. Processed data for genomic sequencing (e.g. CRISPR, targeted amplicon sequencing, and RNA sequencing) and other forms of high-throughput experiments are provided as processed quantifications in Supplementary Datasets. Genomic sequencing raw data are being deposited to Gene Expression Omnibus (GEO) with accession number GSE257805. All data and materials that support the findings of this research are available from the corresponding author upon reasonable request to the academic community.

Temporary Reviewer Token of GEO data:

To review GEO accession GSE257805:

Go to https://nam12.safelinks.protection.outlook.com/?url=https%3A%2F%2Fwww.ncbi.nlm.nih.gov%2Fgeo%2Fquery%2Facc.cgi%3Facc%3DGSE257805&data=05%7C02%7Cchuanpeng.dong%40yale.edu%7Ce32121e485174a38e00608dcc57d1efe%7Cdd8cbebb21394df8b4114e3e87abeb5c%7C0%7C0%7C638602388403905272%7CUnknown%7CTWFpbGZsb3d8eyJWIjoiMC4wLjAwMDAiLCJQIjoiV2luMzIiLCJBTiI6Ik1haWwiLCJXVCI6Mn0%3D%7C0%7C%7C%7C&sdata=M3sjk20x8u%2B6lkXFQaV6GwKWbbW4CQu%2FQvFmCGlwk6s%3D&reserved=0

Enter token slghsgqaxdinfef into the box

### Code availability

Analytic codes used to generate figures that support the findings of this study will be available from the corresponding author upon request.

### Life sciences study design

#### Sample size determination

Sample size was determined according to the lab’s prior work or similar approaches in the field.

#### Data exclusions

No samples were excluded from data analyses.

#### Replication

All experiments were done with at least three biological replicates or two infection replicates. Experimental replications were indicated in detail in methods section and in each figure panel’s legend.

#### Blinding

Investigators were not blinded in *in vitro* experiments. In certain NGS data analysis, such as CRISPR screen and RNA sequencing, investigators were blinded for initial processing of the original data using key-coded metadata.

### Reporting for specific materials, systems and methods

**Antibodies**

|  |  |  |  |
| --- | --- | --- | --- |
| Mouse PD1 | BioXCell | BE0146 | Neturalization antibody |
| Human TIM3 | Biologend | 345009 | Neturalization antibody |
| yH2AX | Cell Signaling Technology | 9718T | Immunofluorescence |
| 53BP1 | Novus Biologicals | NB100-904 | Immunofluorescence |
| RPA32 | Bethyl Laboratories | A300-246A | Immunofluorescence |
| Mouse PDL1-APC | Biologend | 124311 | Flow cytometry |
| OVA-MHCII APC | Biologend | 141605 | Flow cytometry |
| Human CEACAM1-PE | Fisher | 53-0668-41 | Flow cytometry |
| Human CEACAM1-488 | Thermofisher | 53-0668-41 | Flow cytometry |
| Human TIM3 PE-Cy7 | Biologend | 119715 | Flow cytometry |
| Flag PE | Biologend | 637310 | Flow cytometry |
| EGFR PE | Biologend | 352903 | Flow cytometry |
| Human STING-488 | Cell Signaling Technology | 78827S | Flow cytometry |
| Human/mouse phospho-TBK1 | Cell Signaling Technology | 13498S | Flow cytometry |
| Mouse phospho-IRF3 | Cell Signaling Technology | 83611S | Flow cytometry |
| Mouse PD1-APC | Biologend | 135210 | Flow cytometry |
| Mouse LAG3-PE | Biologend | 125207 | Flow cytometry |
| Mouse TIGIT-APC | Biologend | 142105 | Flow cytometry |
| Mouse CD160-PE | Biologend | 143003 | Flow cytometry |
| Mouse CEACAM1-APC | Biologend | 134509 | Flow cytometry |
| Mouse CD45-BV421 | Biologend | 103133 | Flow cytometry |
| Mouse CD45-PE | Biologend | 109807 | Flow cytometry |
| Mouse CD3-PE/Cy7 | Biologend | 100219 | Flow cytometry |
| Mouse CD4-FITC | Biologend | 100509 | Flow cytometry |
| Mouse CD8a-BV605 | Biologend | 100743 | Flow cytometry |
| Mouse CD279 (PD1)-APC | Biologend | 135209 | Flow cytometry |
| Mouse CD223 (LAG3)-PE | Biologend | 125207 | Flow cytometry |
| Mouse MHCII-FITC | Biologend | 107605 | Flow cytometry |
| Mouse CD11b-BV421 | Biologend | 101235 | Flow cytometry |
| Mouse F4/80-PE/Cy7 | Biologend | 123113 | Flow cytometry |
| Mouse CD103-BV605 | Biologend | 121433 | Flow cytometry |
| Mouse CD11c-APC | Biologend | 117309 | Flow cytometry |
| Mouse Capza3 | Santa Cruz | sc-130480 | Western Blot |
| Human TIM3 PE-Cy7 | Biologend | 119715 | Flow cytometry |

**Plasmids**

1 Plasmids
|  |  |  |  |
| --- | --- | --- | --- |
| Brie genome-scale CRISPR library | Addgene | 73633 | CRISPR lentiviral library |
| lentiCas9-Blast | Addgene | 52962 | Generate B16F10-Cas9 line |
| OVA-mCherry | Chen lab | NA | Generate B16F10-Cas9-OVA line to be used in screen |
| LRG (Lenti_sgRNA_EFS_GFP) | Addgene | 65656 | Generage *Capza3* KO in B16F10 |
| H2B-mCherry | Addgene | 20972 | Express H2B-mCherry in B16F10 *Capza3* KO |
| HR extra-chromosomal luciferase reporter | Glazer lab | NA | Assess to HR rate in B16F10, MDA231, PANC1, SKMEL28 |
| NHEJ extra-chromosomal luciferase reporter | Glazer lab | NA | Assess to HR rate in B16F10, MDA231, PANC1, SKMEL28 |
| Luciferase transfection control | Glazer lab | NA | Assess to HR rate in B16F10, MDA231, PANC1, SKMEL28 |
| pU6-(BbsI)_CBh-Cas9-T2A-BFP-P2A-Ad4R1B | Addgene | 64218 | Generate PANC1 AAVS1 TLR line |
| pU6-sgRosa26-1_CBh-Cas9-T2A-BFP | Addgene | 64216 | Generate PANC1 AAVS1 TLR line |
| pAAVS1-TLR targeting vector | Addgene | 64215 | Generate PANC1 AAVS1 TLR line |

### Antibody validation

Antibodies were validated based on manufacturing instructions.

### Eukaryotic cell lines

**Cell line source(s)**

7 Cell line source(s)
|  |  |  |
| --- | --- | --- |
| B16F10 | ATCC | CRL-6475 |
| MDA231 | ATCC | CRM-HTB-26 |
| PANC1 | ATCC | CRL-1469 |
| Human PBMC | StemCell Technologies | 70025 |

### Authentication

Cell lines were authenticated by the commercial vendor.

### Mycoplasma contamination

All the cell lines used here tested negative for mycoplasma contamination.

### Commonly misidentified lines (See ICLAC register)

No commonly misidentified line was used in the study.

### Animals and other organisms

#### Laboratory animals

Mice of both sexes, between age 8 and 12 weeks, were used for the study. 8-week-old OT-I mice were purchased from the Jackson Laboratory and bred in-house. 8-week-old C57BL/6Nr mice were purchase from Charles River lab. 8-week-old NCr mice were purchase from Charles River lab.

#### Wild animals

N/A

#### Field-collected samples

N/A

## Supplementary Tables

**Table S1**.

Oligo sequences used in this study listed in a table.

## Supplementary Datasets

**Dataset S1.**

CRISPR screen processed data and analyses.

**Dataset S2**

RNA-seq processed data and analyses.

**Dataset S3**

Source data and statistics of non-NGS type data provided in an excel file.

**Dataset S4**

TCGA analyses

**Dataset S5**

ICI patient datasets.

## Supplementary Images

**Supplementary Image Set 1**

Raw images for all immunofluorescence pictures used in the manuscript and S3d western blot of CAPZA3.

## Codes

Zip of codes for NGS data analysis

## Supplementary Figure Legend

**Supplementary Figure 1.**
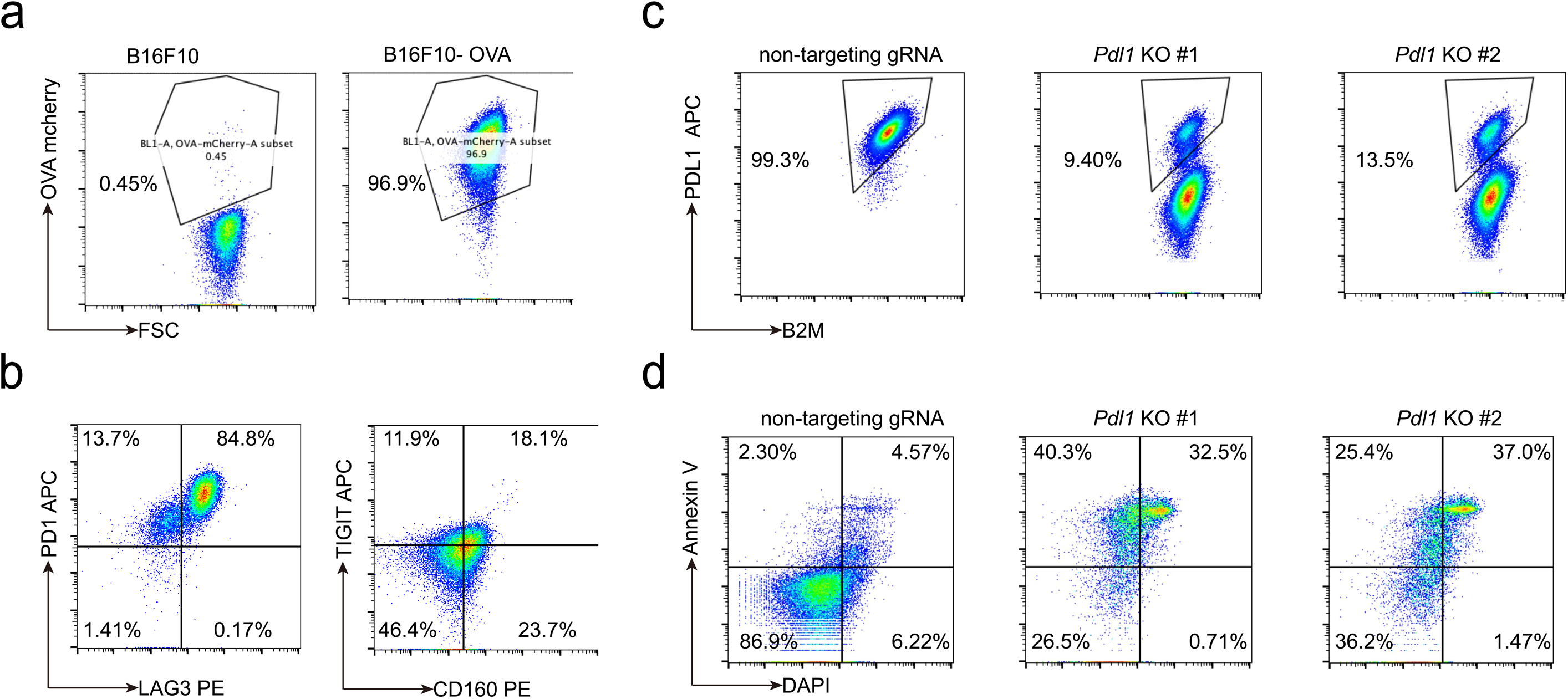
Validation of the screening method using co-culture of *Pdl1* KO B16F10 cells expressing Cas9 protein and OVA antigen (B16F10) with OT-I CD8 T cells. **a,** Flow analysis of OVA antigen expression in Cas9 expressing B16F10 cells used for the CRISPR screen. **b,** Flow analysis of PD1, LAG3, TIGIT and CD160 expression in OT-I CD8 T cells. Prior to co-culture with B16F10-OVA cells, CD8 T cells were isolated from OT-I mice and expanded with OVA antigen and cytokines IL-7 and IL-15 for 3 days. **c,** Flow cytometry analysis of PDL1 in B16F10 cells transduced with lentivirus containing either vector with non-targeting gRNA (WT control) or a vector carrying either *Pdl1* targeting gRNA #1 (PDL1 KO #1) or *Pdl1* targeting gRNA #2 (PDL1 KO #2). **d,** Annexin V and DAPI staining of control and *Pdl1* KO B16F10-OVA cells following co-culture with OT-I CD8 T cells. T:E ratio of 1:1.

**Supplementary Figure 2.**
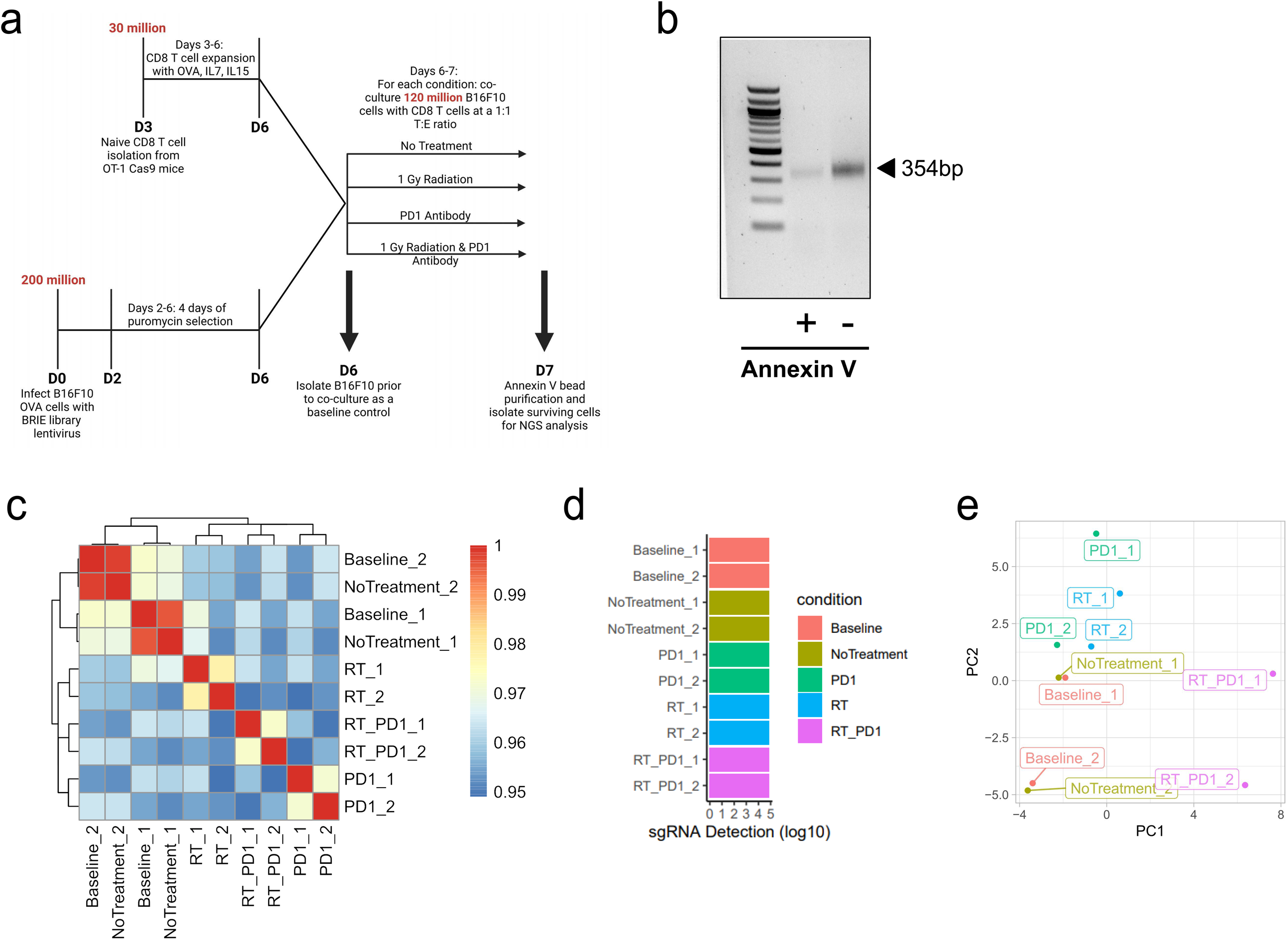
Sequencing analysis of enriched gRNAs in surviving B16F10-OVA cells following co-culture with OT-I CD8 T cells in genome scale CRISPR screens. **a,** Schematic of B16F10 CRISPR screen. Total B16F10 cell number at each step is shown. 2 independent replicates of each of the 4 treatment conditions were done and analyzed for enriched gRNAs. **b,** PCR amplification of gRNA barcode regions in Annexin V positive and Annexin V negative B16F10 fractions from genomic DNA of Annexin V column purified B16F10 cells. **c,** Correlation between 2 independent replicates of the screen. Baseline refers to control B16F10 cells that were isolated following lentiviral infection but prior to any treatment (with radiation or anti-PD1 antibody) and prior to co-culture with CD8 T cells. RT refers to radiation treatment alone, PD1 refers to anti-PD1 antibody treatment alone and RT_PD1 refers to the combination of radiation and anti-PD1 antibody treatment. 1 or 2 within the labels refers to replicate 1 or replicate 2. **d,** sgRNA detection for baseline and each of the treatment conditions (no treatment, radiation alone, anti-PD1 antibody, radiation and anti-PD1 antibody). 1 or 2 within the labels refers to replicate 1 or replicate 2. **e,** PCA analysis of different treatment conditions between the two independent replicates of the screen.

**Supplementary Figure 3.**
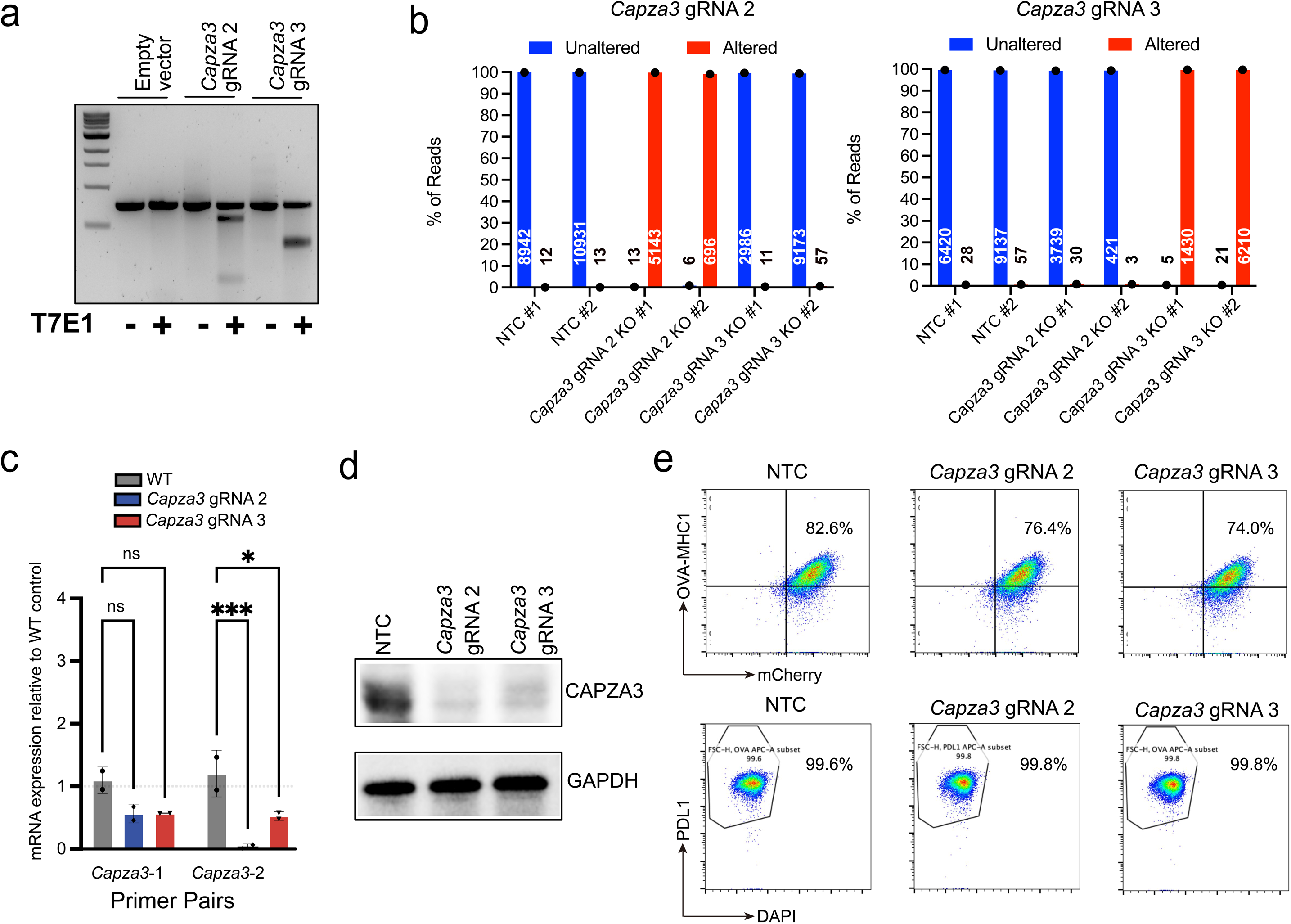
Generation of *Capza3* KO B16F10 cell lines. **a,** T7 endonuclease I (T7E1) analysis of the 2 *Capza3* targeting gRNAs (gRNA 2 and gRNA 3) that were significantly enriched in surviving B16F10 cells from CRISPR genome-scale screens. **b,** Next generation sequencing results for B16F10 for WT and *CAPZA3* KOs are shown. Graphs show the percent of reads with altered sequences within the *CAPZA3* gRNA 2 and *CAPZA3* gRNA 3 target sequences. The number of reads is shown within or above each bar. **c,** qPCR analysis of *Capza3* expression in B16F10 cell lines transduced with lentivirus containing the 2 *Capza3* targeting gRNAs (gRNA 2 and gRNA 3) that were significantly enriched in surviving B16F10 cells from CRISPR genome-scale screens. Significance testing was performed with two-way ANOVA, two biological replicates were done for each sample, * p < 0.05, ** p< 0.01, *** p < 0.001, **** p < 0.0001. **d,** Western blot of CAPZA3 in WT and *Capza3* KO B16F10 cell lines. **e,** Analysis of OVA expression (top) and PDL1 expression (bottom) in B16F10 cells transduced with either lentivirus containing a non-targeting control gRNA or the 2 *Capza3* targeting gRNAs that were enriched in the CRISPR genome-scale screens.

**Supplementary Figure 4.**
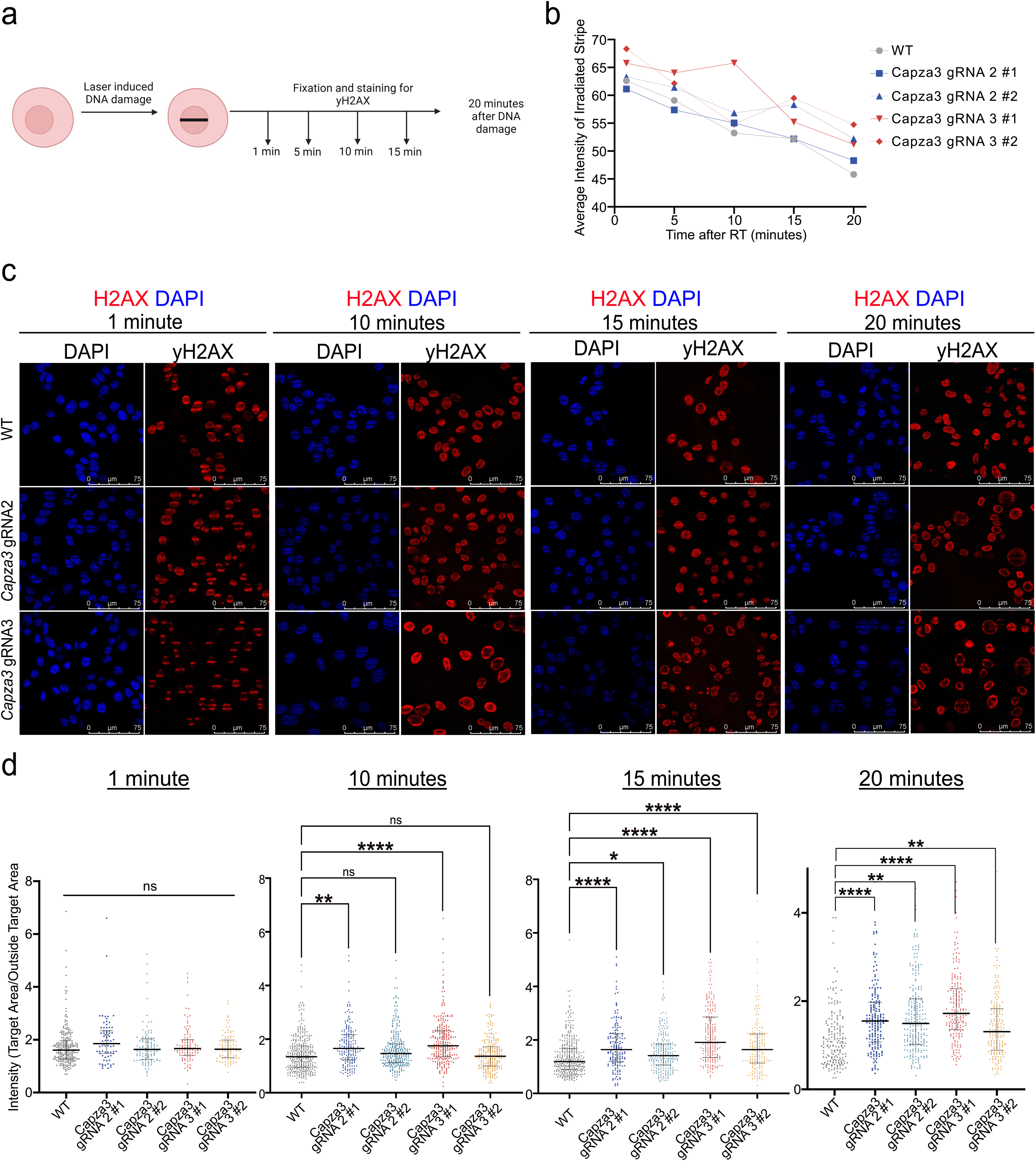
Laser micro-irradiation induced DNA damage and staining with γH2AX in WT and *Capza3* KO cells. **a,** Schematic of laser micro-irradiation targeted subnuclear DNA damage followed by fixation and staining with γH2AX at different time points. **b,** Average intensity of γH2AX stripe in the region of laser induced DNA damage in WT and *Capza3* KO cells at different time points after laser micro-irradiation. Significance testing was performed with two-way ANOVA, the average intensity from three biological replicates is shown. P-values: WT vs. *Capza3* gRNA 2 KO #1: ns, WT vs *Capza3* gRNA 2 KO #2: ns, WT vs *Capza3* gRNA 3 KO #1: 0.0082, WT vs *Capza3* gRNA 3 KO #2: 0.0157. **c,** Immunofluorescence imaging of γH2AX DNA damage foci 1 minute, 10 minutes, 15 minutes, and 20 minutes following laser micro-irradiation in WT and *Capza3* KO cells. **d,** Quantification in individual cells of the ratio of intensity of γH2AX staining in the region of laser induced DNA damage versus outside this region at 1 minute, 10 minutes, 15 minutes and 20 minutes after laser micro-irradiation. Significance testing was performed with two-way ANOVA, individual cell data from three biological replicates were pooled together for each sample. For all experiments, WT cells were transduced with a lentiviral vector expressing a NTC gRNA. * p < 0.05, ** p < 0.01, *** p < 0.001, **** p < 0.0001.

**Supplementary Figure 5.**
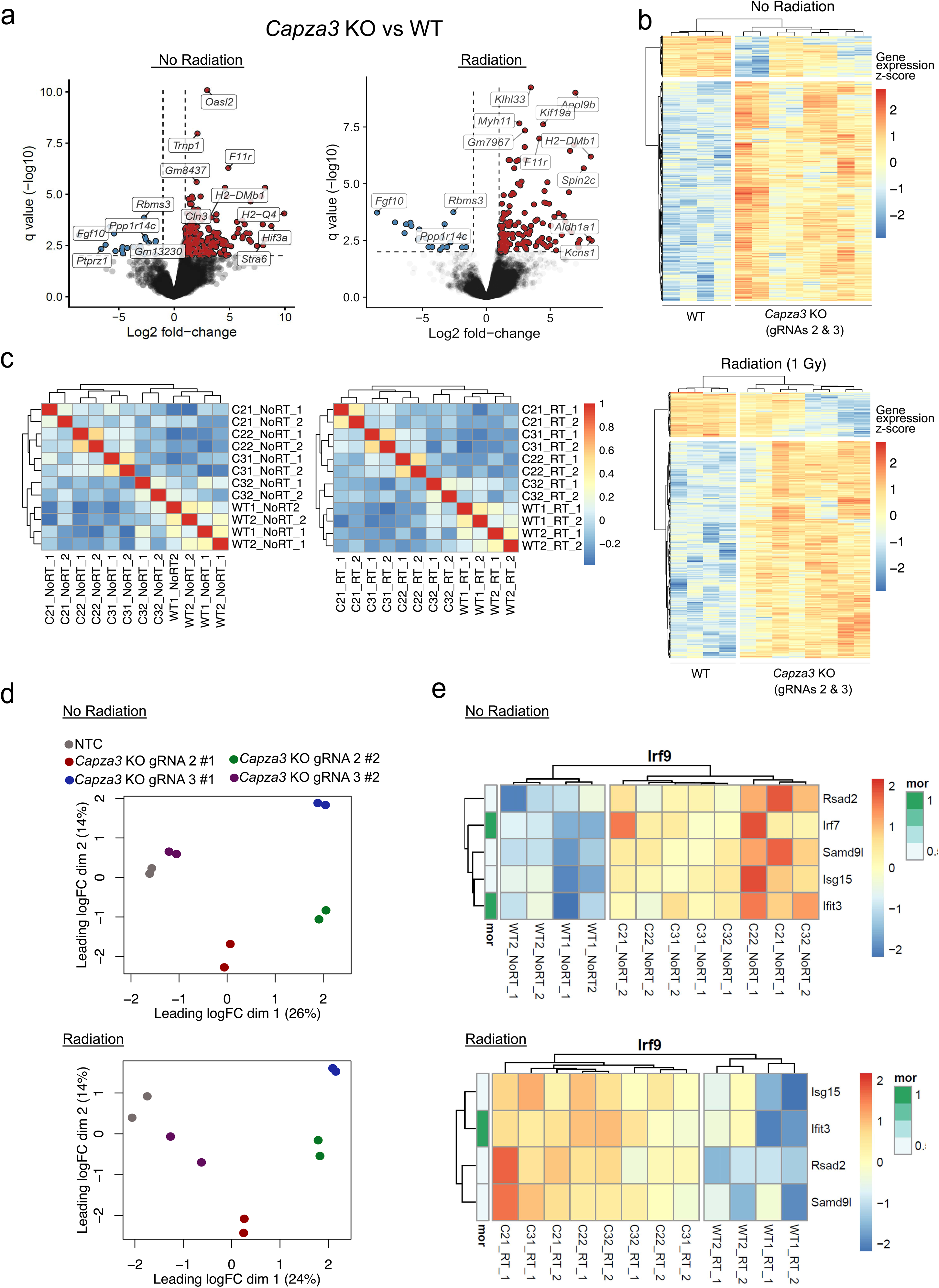
RNA sequencing analysis of B16F10 WT and *Capza3* gRNA 2 and *Capza3* gRNA 3 KO cells before and after 1 Gy radiation treatment. **a,** Volcano plot of genes differentially expressed between *Capza3* KO and WT cells before radiation treatment (left) and after radiation treatment (right) from bulk RNA-Seq analysis of WT and *Capza3* KO B16F10 cells. **b,** Heat map showing differentially expressed genes between WT and *Capza3* KO B16F10 cells before (top) and after (bottom) low-dose (1 Gy) radiation treatment from bulk RNA-Seq analysis of WT and *Capza3* KO B16F10 cells. **c,** Correlation between 2 independent replicates of RNA sequencing analysis. C21_1 refers to *Capza3* KO gRNA 2, clone 1 and replicate 1, C32_2 refers *Capza3* KO gRNA 3, clone 2 and replicate 2, etc. All samples before radiation treatment are shown on the left (12 samples in total) and all samples after radiation treatment are shown on the right (12 samples total). **d,** PCA analysis of WT and *Capza3* KO mutants from 2 independent replicates of RNA sequencing analysis. All samples before radiation treatment are shown on the top panel and all samples after radiation treatment are shown on the bottom panel. **e,** Targets of *Irf9* transcription factor that show the greatest upregulation in *Capza3* KO compared to WT B16F10 cells, before radiation (top panel) and after radiation (bottom panel). For RNA-Seq analyses, n = 2 replicates for each cell line (WT #1, WT #2, *Capza3* gRNA 2 KO #1, *Capza3* gRNA 2 KO #2, *Capza3* gRNA 3 KO #1, *Capza3* gRNA 3 KO #2) and each condition (no radiation or after 1 Gy radiation) were done, for 24 samples in total.

**Supplementary Figure 6.**
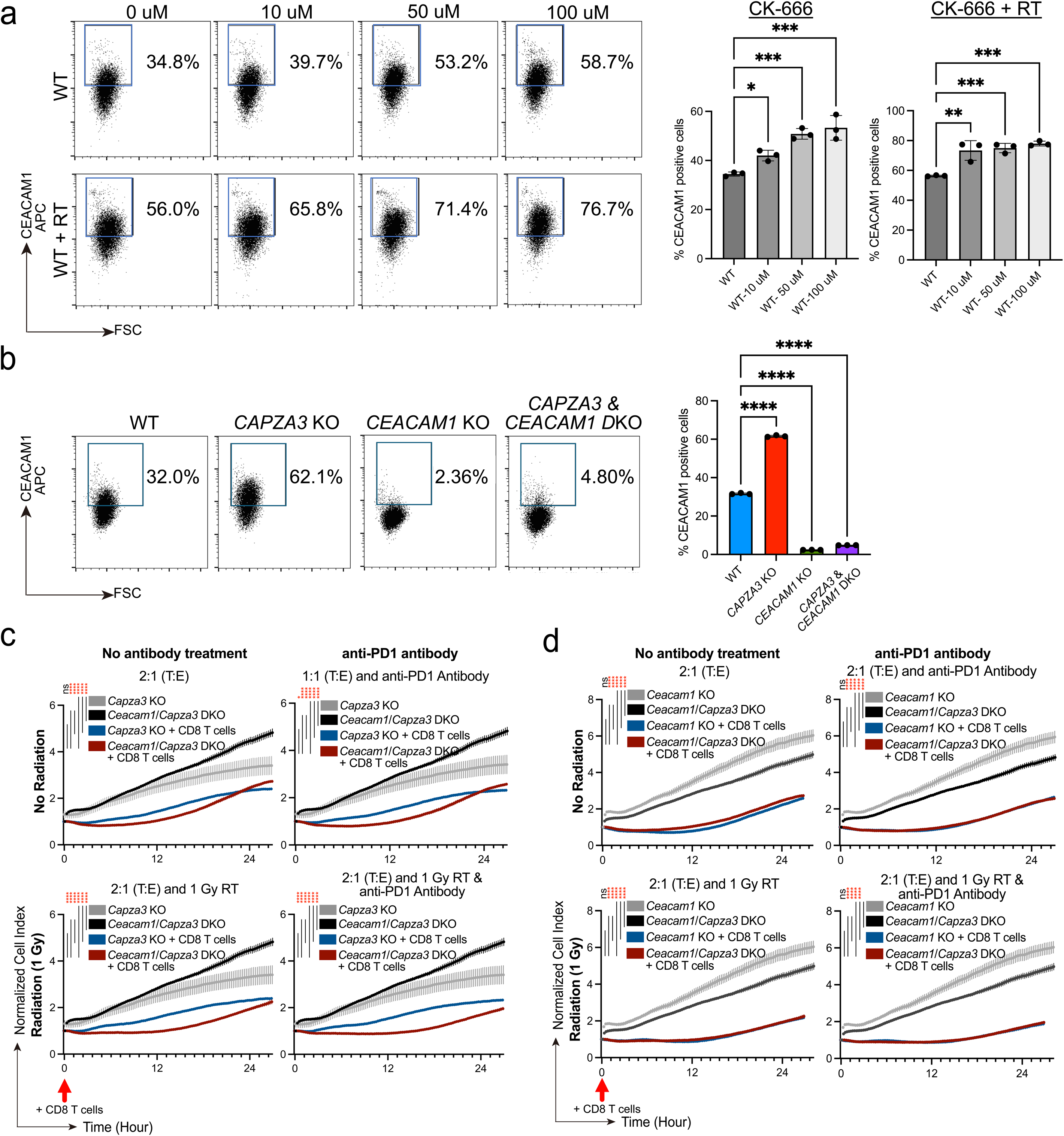
Expression of CEACAM1 following CK-666 treatment and generation of *Ceacam1* KO and *Capza3*/*Ceacam1* DKO B16F10 cells. **a,** Flow analysis of CEACAM1 positive cells following treatment of WT B16F10 cells with CK-666 at different concentrations with and without exposure to radiation. Representative flow plots are shown on the left and quantification is shown on right. Significance testing was performed with one-way ANOVA. A minimum of 3 biological replicates were done. **b,** Flow analysis of CEACAM1 positive cells in WT, *Capza3* KO, *Ceacam1* KO and *Capza3*/*Ceacam1* DKO cell lines. Significance testing was performed with one-way ANOVA. A minimum of 3 biological replicates were done. **c,** Cell attachment index of *Capza3* KO B16F10 and *Capza3*/*Ceacam1* DKO B16F10 cells co-cultured with OT-1 T cells. Cells were either just exposed to OT-1 CD8 T cells (2:1 T:E) or exposed to CD8 T cells with: 1 Gy low-dose radiation (2:1 T:E and 1 Gy RT), anti-PD1 antibody (2:1 T:E and anti-PD1 Antibody) or 1 Gy low-dose radiation and anti-PD1 antibody (2:1 T:E and 1 Gy RT + anti-PD1 Antibody). Experiments used a 2:1 T:E (tumor: effector) ratio. Time point of addition of OT-1 CD8 T cells is shown by an arrow. Data shown as means ± SEM. *P* value was determined by two-way ANOVA. Data are representative of three biological replicates. **d,** Cell attachment index of *Ceacam1* KO B16F10 and *Capza3*/*Ceacam1* DKO B16F10 cells co-cultured with OT-1 T cells. Cells were either just exposed to OT-1 CD8 T cells (2:1 T:E) or exposed to CD8 T cells with: 1 Gy low-dose radiation (2:1 T:E and 1 Gy RT), anti-PD1 antibody (2:1 T:E and anti-PD1 Antibody) or 1 Gy low-dose radiation and anti-PD1 antibody (2:1 T:E and 1 Gy RT + anti-PD1 Antibody). Experiments used a 2:1 T:E (tumor: effector) ratio. Time point of addition of OT-1 CD8 T cells is shown by an arrow. Data shown as means ± SEM. *P* value was determined by two-way ANOVA. Data are representative of three biological replicates. * p < 0.05, ** p < 0.01, *** p < 0.001, **** p < 0.0001.

**Supplementary Figure 7.**
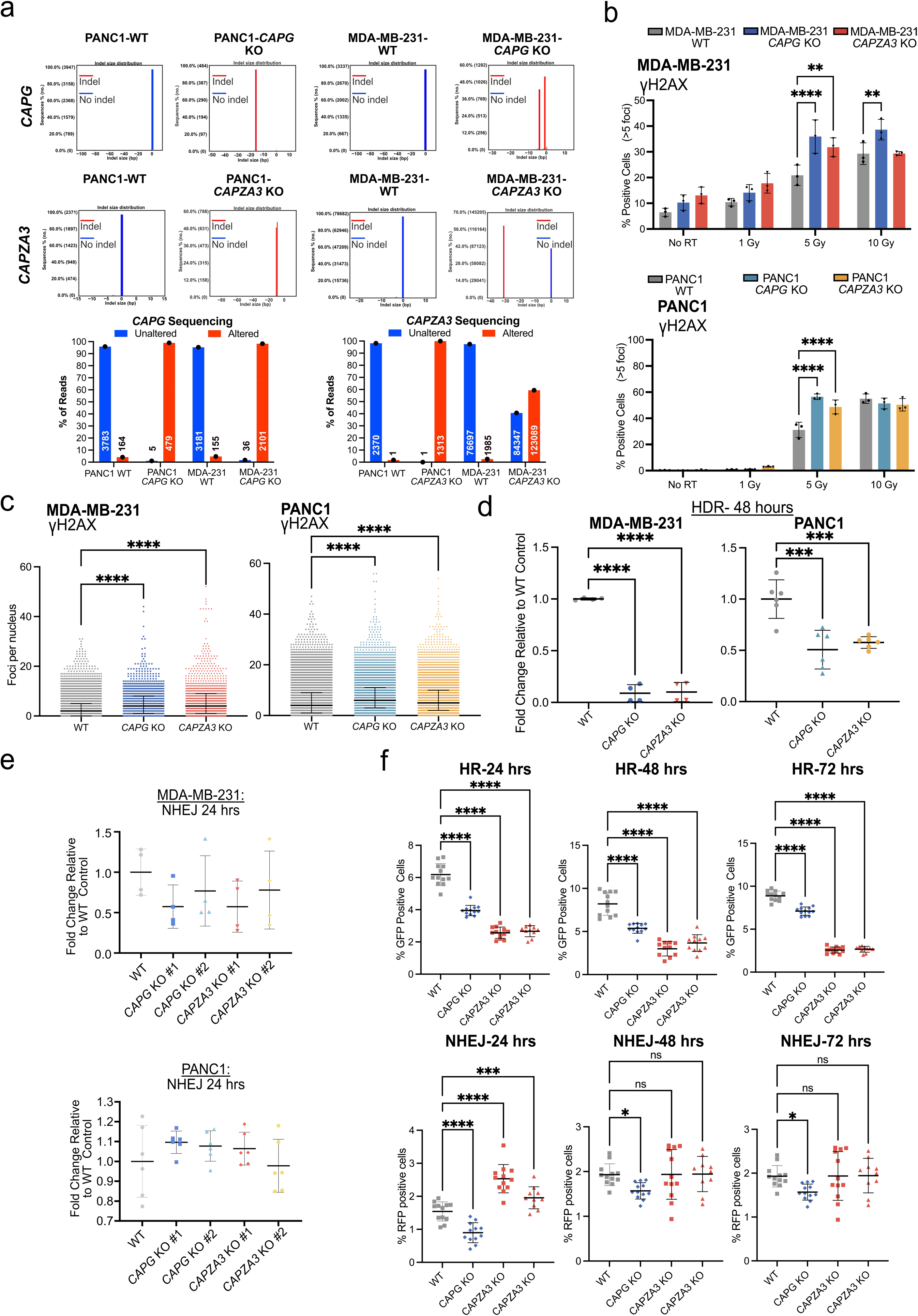
Analysis of DNA damage foci and repair efficiency in PANC1 and MDA-MB-231 *CAPG* and *CAPZA3* KO cell lines. **a,** Next generation sequencing results for PANC1 and MDA-MB-231 *CAPG* and *CAPZA3* KOs are shown. The top panel shows the percent of sequencing reads with indel mutations, with the size of the indel mutation shown on the x axis. The bottom graphs show the percent of reads with altered sequences within the *CAPG* or *CAPZA3* gRNA target sequences. The number of reads is shown within or above each bar. **b,** Quantification of γH2AX foci in WT and additional clonal *CAPG* and *CAPZA3* KO MDA-MB-231 (left) and PANC1 (right) cell lines 24 hours after different doses of radiation treatment. Significance testing was performed with two-way ANOVA, three biological replicates were done for each sample and treatment condition. **c,** Quantification of total γH2AX foci in individual cells in WT and additional clonal *CAPG* and *CAPZA3* KO MDA-MB-231 (left) and PANC1 (right) cell lines 24 hours after exposure to 5 Gy radiation treatment. Significance testing was performed with one-way ANOVA, individual cell data from three biological replicates were pooled together for each sample. P-values are shown for comparisons of the KO line (*CAPG* KO or *CAPZA3* KO) to WT. **d,** Quantification of luciferase activity, compared to WT cells, 48 hours after transfection with a linearized HDR extrachromosomal reporter in additional clonal *CAPG* and *CAPZA3* KO MDA-MB-231 (left) and PANC1 (right) cell lines. Significance testing was performed with one-way ANOVA. 4 biological replicates were done for the MDA-MB-231 and 6 biological replicates were done for the PANC1. **e,** Quantification of luciferase activity, compared to WT cells, 48 hours after transfection with a linearized NHEJ extrachromosomal reporter in clonal *CAPG* and *CAPZA3* KO MDA-MB-231 (top) and PANC1 (bottom) cell lines. Significance testing was performed with one-way ANOVA. 4 biological replicates were done for the MDA-MB-231 and 6 biological replicates were done for PANC1. **f,** Analysis of additional *CAPG* and *CAPZA3* KO PANC1 cell lines with the AAVS1 TLR reporter. Quantification of GFP positive cells (top panel, % cells with successful HDR) and RFP positive cells (bottom panel, % cells with successful NHEJ) 24, 48 and 72 hours after transfection of *Rosa26* gRNA in PANC1 *AAVS1* TLR WT, *CAPG* and *CAPZA3* KO cell lines. Significance testing was performed with two-way ANOVA, four biological replicates were done for each sample. For all experiments, WT cells were passage-matched controls that underwent the procedure to generate KO cell lines (transfected with plasmid containing *CAPG* or *CAPZA3* targeting gRNA, sorted as single cells into a 96 well plate and expanded) but did not have a mutation at either the *CAPG* or *CAPZA3* locus. * p < 0.05, ** p< 0.01, *** p < 0.001, **** p < 0.0001.

**Supplementary Figure 8.**
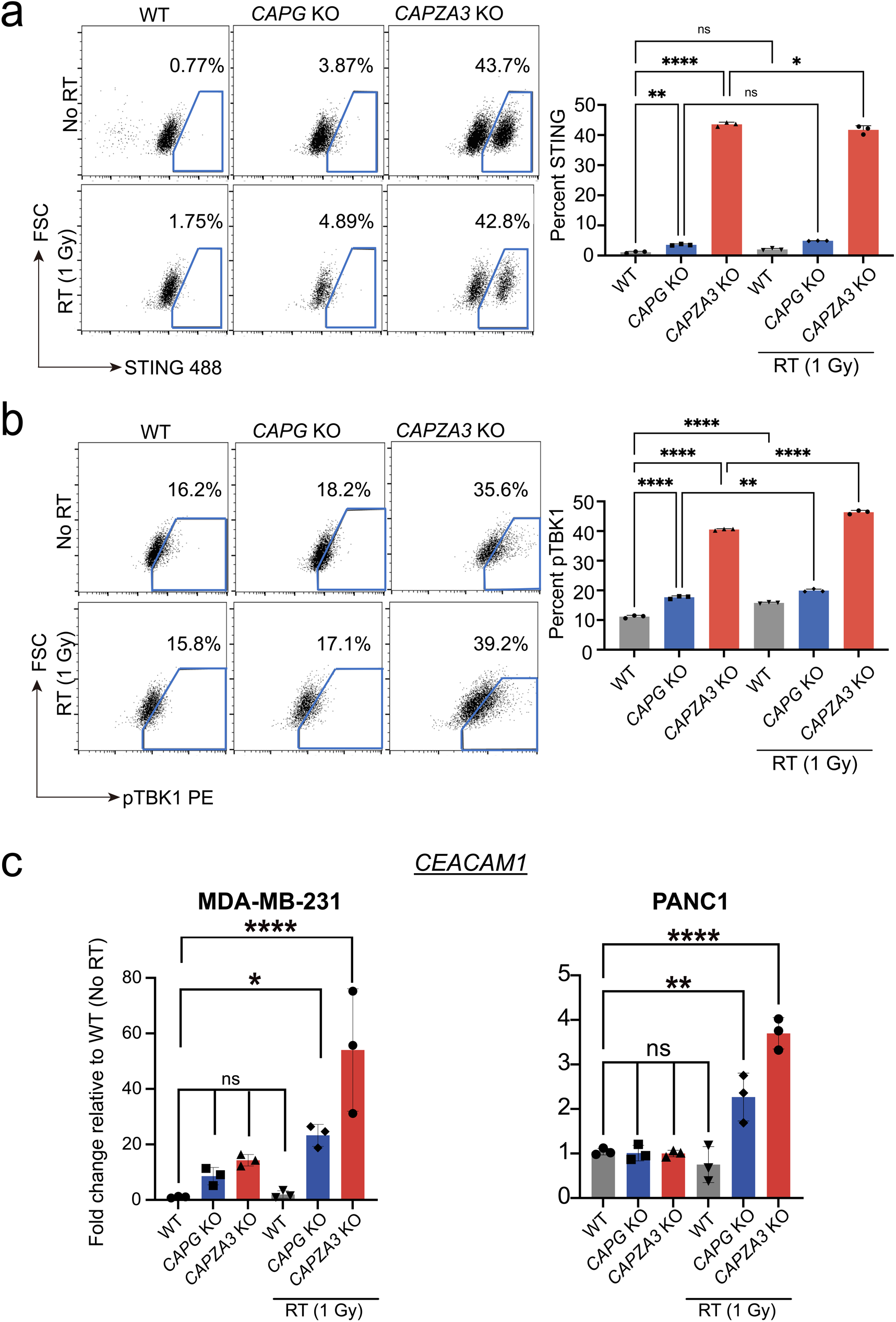
Activation of the STING pathway and expression of *CEACAM1* in MDA-MB-231 and PANC1 cell lines after radiation treatment. **a,** Representative flow plots of STING staining in WT, *CAPG* and *CAPZA3* KO MDA-MB-231 cell lines before and after low-dose (1 Gy) radiation treatment. Representative plots are showed on the left and quantification of positive cells is shown on the right. Significance testing was performed with two-way ANOVA, three biological replicates were done for each sample and treatment condition. **b,** Representative flow plots of phospho-TBK1 staining in WT, *CAPG* and *CAPZA3* KO MDA-MB-231 cell lines before and after low-dose (1 Gy) radiation treatment. Representative plots are showed on the left and quantification of positive cells is shown on the right. Significance testing was performed with two-way ANOVA, three biological replicates were done for each sample and treatment condition. **c,** qPCR analysis of *CEACAM1* expression in MDA-MB-231 (left) and PANC1 (right) WT and KO cell lines before and after low-dose (1 Gy) radiation treatment. Significance testing was performed with two-way ANOVA, three biological replicates were done for each sample and treatment condition.

**Supplementary Figure 9.**
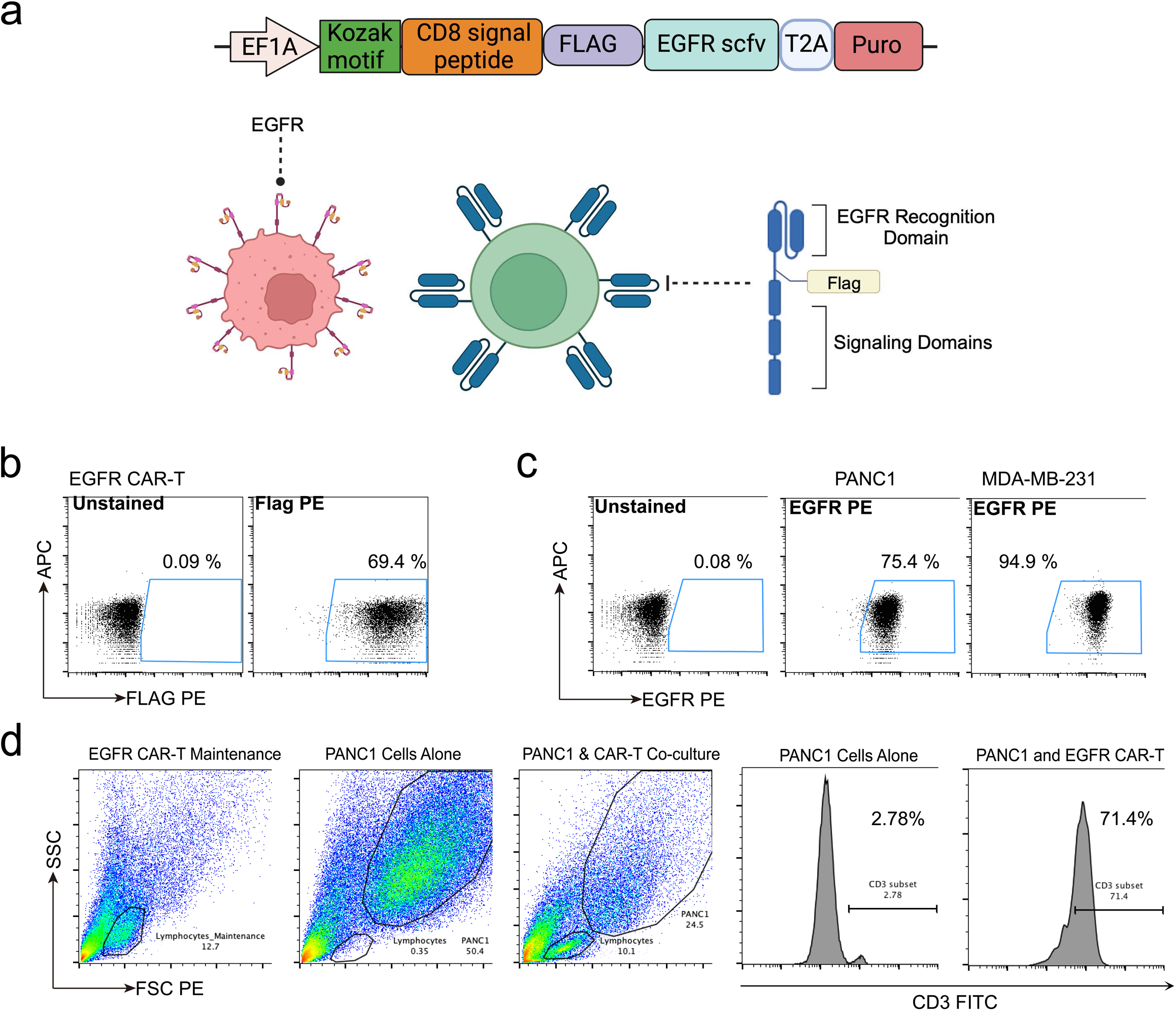
Cell survival of MDA-MB-231 *CAPG* and *CAPZA3* KO cell lines compared to WT cells following treatment with radiation, anti-TIM3 antibody or anti-PD1 antibody and co-culture with EGFR CAR-T. **a,** Schematic of EGFR CAR-T construct. **b,** Representative flow plots of EGFR chimeric antigen receptor (CAR) Flag tag on EGFR CAR-T cells. **c,** Representative flow plots of EGFR expression in MDA-MB-231 and PANC1 cell lines. **d,** Gating strategy to quantify surviving cancer cells following co-culture with EGFR CAR-T is shown. Cancer cells and lymphocytes were first gated based on forward and side scatter properties and then lymphocytes were identified using CD3-FITC staining.

**Supplementary Figure 10.**
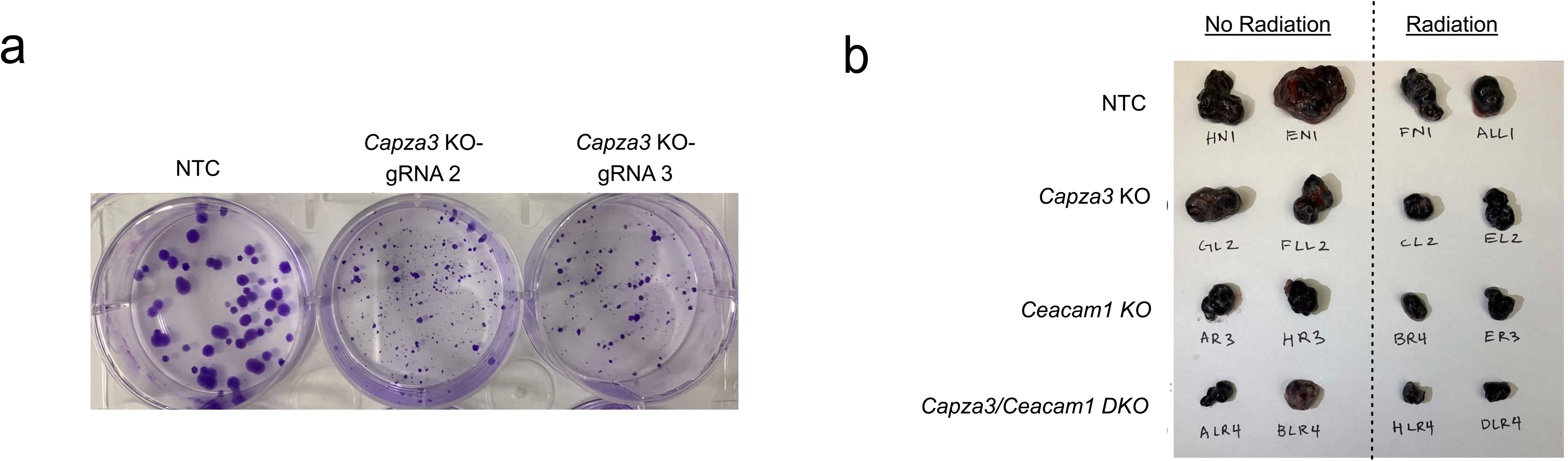
*In-vivo* studies of WT, *Capza3* KO, *Ceacam1* KO and *Capza3*/*Ceacam1* DKO B16F10 cells in C57BL/6 mice. **a,** Representative images of in-vitro colony formation for WT and *Capza3* KO B16F10 cells. **b,** Representative gross tumors harvested from WT, *Capza3* KO, *Ceacam1* KO and *Capza3*/*Ceacam1* DKO B16F10 cells with and without radiation treatment.

**Supplementary Figure 11.**
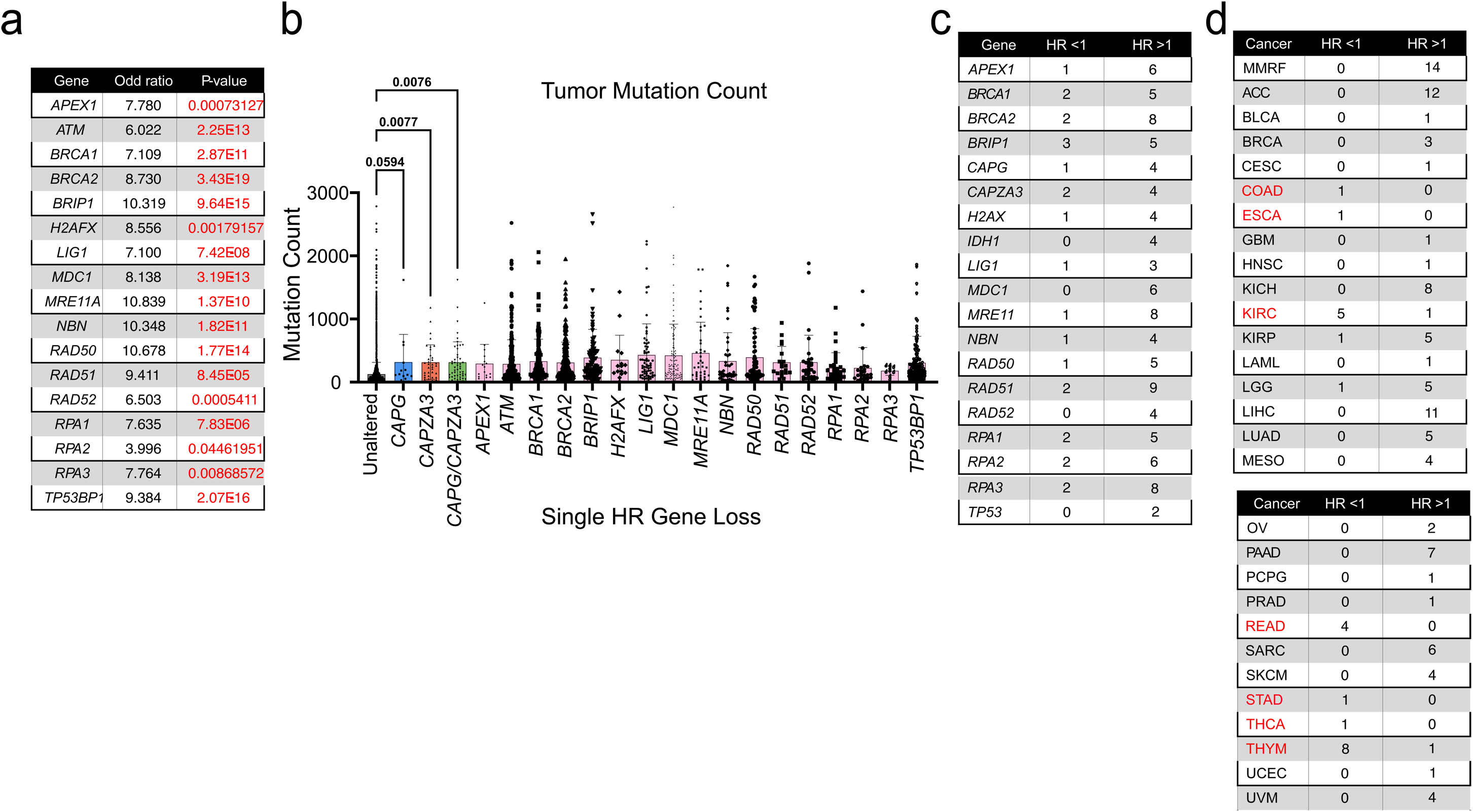
TCGA analysis of tumor mutation count associated with inactivating mutations in *CAPG* and *CAPZA3* and correlation of *CAPG* and *CAPZA3* expression with overall survival for different cancer types. **a,** Analysis of the statistical significance of the co-occurrence of *CAPG*/*CAPZA3* mutations with other HDR genes mutations evaluated using Fisher’s exact test. **b,** Analysis of tumor mutation count in patients with an isolated inactivating mutation in *CAPG* or *CAPZA3* or a mutation in a single HDR associated gene. P-values are shown for selected comparisons. Significance testing was performed with one-way ANOVA. **c,** The number of cancers for which a HDR gene was prognostic for patient OS are shown. The number of cancers in which high expression of the HDR gene was associated with poorer OS are shown (HR > 1). The number of cancers in which high expression of the HDR gene was associated with improved OS (HR < 1) are shown. **d,** The number of HDR genes that were prognostic for patient OS for each cancer type are shown. The number of HDR genes for which high expression was associated with poorer OS in each cancer are shown (HR > 1). The number of HDR genes for which high expression of the HDR gene was associated with improved OS in each cancer (HR < 1) are shown. Cancers in which high expression of an HDR gene was associated with improved survival (or low expression was associated with poorer survival) are highlighted in red.

## Notes

### Competing Interest Statement

The authors have declared no competing interest.

### Summary of Updates

Added new data, revised most figures, updated author list.

